# Rapid host switching of *Wolbachia* and even more rapid turnover of their phages and incompatibility-causing loci

**DOI:** 10.1101/2023.12.04.569981

**Authors:** J. Dylan Shropshire, William R. Conner, Dan Vanderpool, Ary A. Hoffmann, Michael Turelli, Brandon S. Cooper

## Abstract

About half of all insect species carry maternally inherited *Wolbachia* alphaproteobacteria, making *Wolbachia* the most common endosymbionts known in nature. Often *Wolbachia* spread to high frequencies within populations due to cytoplasmic incompatibility (CI), a *Wolbachia*-induced sperm modification caused by prophage-associated genes (*cifs*) that kill embryos without *Wolbachia*. Several *Wolbachia* variants also block viruses, including *w*Mel from *Drosophila melanogaster* when transinfected into the mosquito *Aedes aegypti*. CI enables the establishment and stable maintenance of pathogen-blocking *w*Mel in natural *Ae. aegypti* populations. These transinfections are reducing dengue disease incidence on multiple continents. While it has long been known that closely related *Wolbachia* occupy distantly related hosts, the timing of *Wolbachia* host switching and molecular evolution has not been widely quantified. We provide a new, conservative calibration for *Wolbachia* chronograms based on examples of co-divergence of *Wolbachia* and their insect hosts. Synthesizing publicly available and new genomic data, we use our calibration to demonstrate that *w*Mel-like variants separated by only about 370,000 years have naturally colonized holometabolous dipteran and hymenopteran insects that diverged approximately 350 million years ago. Data from *Wolbachia* variants closely related to those currently dominant in *D. melanogaster* and *D. simulans* illustrate that *cifs* are rapidly acquired and lost among *Wolbachia* genomes, on a time scale of 10^4^–10^5^ years. This turnover occurs with and without the *Wovirus* prophages that contain them, with closely related *cifs* found in distantly related phages and distantly related *cifs* found in closely related phages. We present evidence for purifying selection on CI rescue function and on particular Cif protein domains. Our results quantify the tempo and mode of rapid host switching and horizontal gene transfer that underlie the spread and diversity of *Wolbachia* sampled from diverse host species. The *w*Mel variants we highlight from hosts in different climates may offer new options for broadening *Wolbachia*-based biocontrol of diseases and pests.

## Introduction

Maternally transmitted *Wolbachia* bacteria were first discovered in the ovaries of the mosquito *Culex pipiens* (Hertig and Wolbach 1924). They are now recognized as the most common endosymbionts in nature, occurring in about half of all insect species (Weinert *et al*. 2015). This includes distantly related host species carrying closely related *Wolbachia* (O’Neill *et al*. 1992), indicative of horizontal *Wolbachia* movement among hosts. *Wolbachia* are known for their effects on host reproduction, with cytoplasmic incompatibility (CI) observed in 10 arthropod host orders (Shropshire *et al*. 2020). CI kills *Wolbachia*-free embryos fertilized by *Wolbachia*-carrying males, often driving the endosymbiont to high frequencies in natural populations (Yen and Barr 1973; Hoffmann *et al*. 1990). CI also enables successful biocontrol of human diseases by facilitating the establishment in vector populations of pathogen-blocking *Wolbachia* like *w*Mel from *Drosophila melanogaster* (Walker *et al*. 2011; Utarini *et al*. 2021; Lenharo 2023; Velez *et al*. 2023). Using published and new data, our goal is to elucidate the timescale of *Wolbachia* movements among host species and the rapid turnover and evolution of genes (*cifs*) that can cause (*cifB* and *cifA*) and rescue (*cifA*) CI (LePage *et al*. 2017; Beckmann *et al*. 2017, 2021; Shropshire and Bordenstein 2019).

The first DNA analyses of *Wolbachia* (O’Neill *et al*. 1992), based on partial sequences of 16S rRNA loci, demonstrated that *Wolbachia* and their insect hosts have deeply discordant phylogenies, with similar *Wolbachia* found in distantly related hosts, including Diptera, Lepidoptera, and Coleoptera. The alternative modes of these host transfers and their rapid time scale have been revealed with increasing accuracy as more molecular data are collected. In their pioneering analysis of divergence times, Werren *et al*. (1995) used a “universal molecular clock” for bacteria (Ochman and Wilson 1987) applied to *Wolbachia ftsZ* sequences extracted from flies and wasps. With no differences over 265 synonymous substitution sites within a 937 bp region of *ftsZ* from *Wolbachia* found in the parasitic hymenopteran *Asobara tabida* and its dipteran host *Drosophila simulans* (from Riverside California), Werren *et al*. (1995) estimated a 95% (99%) confidence interval for the divergence time of these *Wolbachia* of 0–1.6 (0–2.5) million years. Recent analyses (Wang *et al*. 2016) suggest that Diptera and Hymenoptera diverged about 350 million years ago (MYA). Many subsequent studies, most recently by Vancaster and Blaxter (2023), have confirmed the pervasive discordance of *Wolbachia* and host phylogenies. In contrast to these generally facultative associations, *Wolbachia* seem to have become obligate symbionts of filarial nematodes (Bandi *et al*. 1998, reviewed in Manoj *et al*. 2021) in which they generally codiverge with their hosts (cf., Moran 2007).

Werren and associates developed a refined chronology of *Wolbachia* movement among host species (Raychoudhury *et al*. 2009). They cross-calibrated *Wolbachia* divergence with host nuclear and mtDNA divergence, exploiting a convincing example of cladogenic transmission (i.e., codivergence) of *Wolbachia* in the wasps *Nasonia longicornis* and *N. giraulti*. The key evidence supporting codivergence was concordant divergence-time estimates for the hosts and *Wolbachia* based on independently derived molecular clocks for synonymous-site divergence of eukaryotic nuclear genes and an updated rate estimate for coding-region divergence across bacteria (Ochman *et al*. 1999). For the host-*Wolbachia* pairs showing plausible cladogenic *Wolbachia* transmission ((*N. longicornis*, *w*NlonB1) and (*N. giraulti*, *w*NgirB)), Raychoudhury *et al*. (2009) estimated that *Wolbachia* diverged at about one-third the rate of the host nuclear genomes for synonymous sites (see our Materials and Methods). In contrast, mtDNA diverged at synonymous sites about 1.2×10^2^ times as fast as co-inherited *Wolbachia.* As summarized in our Materials and Methods, the Raychoudhury *et al*. (2009) data imply an average *Wolbachia* yearly substitution rate at third-codon sites of approximately 2.2×10^−9^.

Richardson *et al*. (2012) produced an alternative approach to calibrating rapid *Wolbachia* divergence, comparing full genomes of *Wolbachia* and mitochondria among *Drosophila melanogaster* lineages. As expected under joint maternal inheritance, *D. melanogaster* isofemale lines produced concordant mtDNA and *Wolbachia* phylogenies. Like Raychoudhury *et al*. (2009), Richardson *et al*. (2012) observed that the third-site mtDNA differences were roughly 10^2^ greater than *Wolbachia* codon differences (which do not differ across the three coding positions over short divergence times, cf. Conner *et al*. 2017, Table 1). Having estimated *relative* sequence divergence for *Wolbachia* and mtDNA among isofemale lines, Richardson *et al*. (2012) estimated an *absolute* rate for *Wolbachia* evolution. Using the per-generation mtDNA mutation rate as a prior in a Bayesian analysis of isofemale-line divergence, they estimated a “short-term” *Wolbachia* third-site substitution rate of 6.87×10^−9^ per site per year (see our Materials and Methods).

**Table 1.**
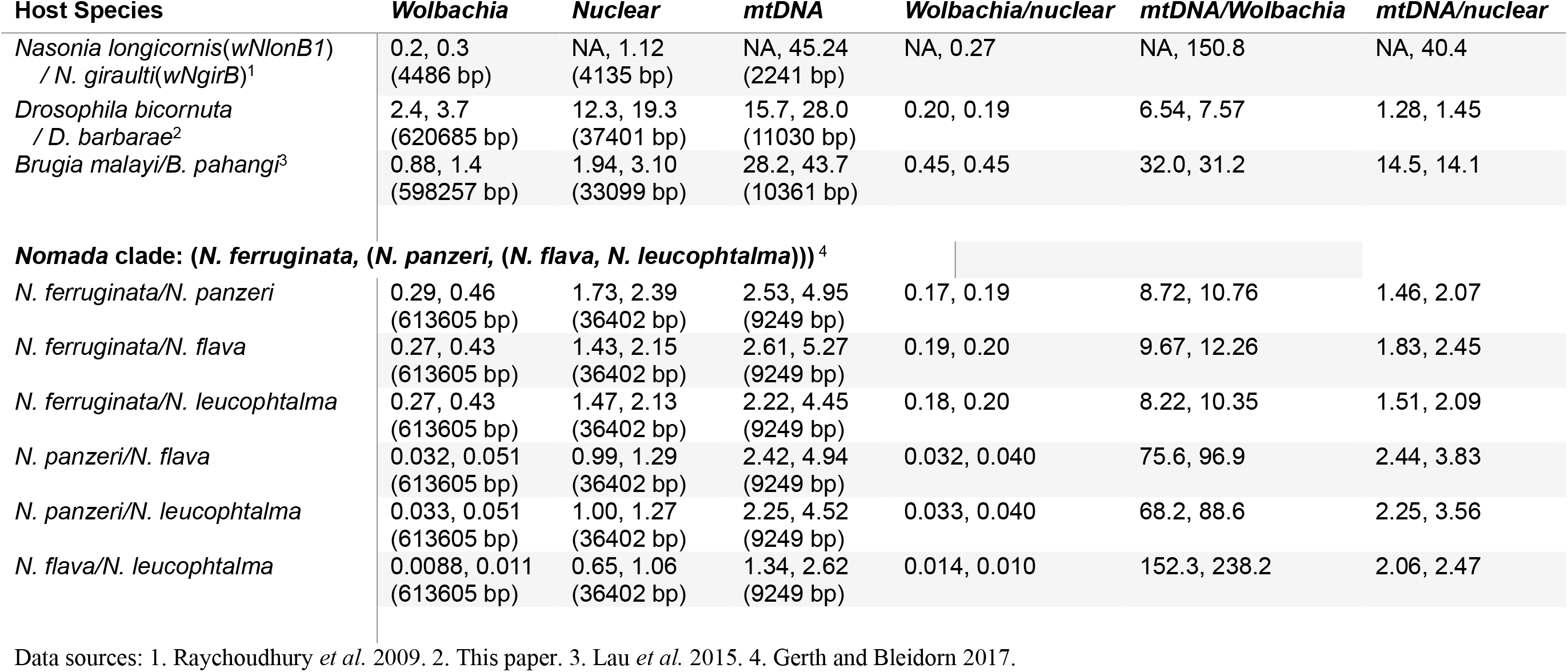
Absolute and relative divergence of *Wolbachia* versus host nuclear and mitochondrial (mtDNA) genomes for cases of putative cladogenic *Wolbachia* transmission. For each entry in the body of the table, the first value is the estimated percent substitutions (after Jukes-Cantor correction) per third-position codon sites, the second value is the estimated percentage of synonymous substitutions across the first and third positions. The values in parentheses are numbers of nucleotides on which the divergence estimates are based.

Turelli *et al*. (2018) applied this short-term calibration of Richardson *et al*. (2012) to estimate divergence times for *Wolbachia* closely related *w*Ri, the first *Wolbachia* variant discovered in a *Drosophila* species (Hoffmann *et al*. 1986). This yielded an estimate that *w*Ri-like strains diverged less than 30 thousand years (KY) occupy *Drosophila* hosts diverged 10–50 million years (MY). Ahmed *et al*. (2016) used a similar calibration to estimate horizontal transmission times for *Wolbachia* among Lepidoptera species. Analyses of model *Drosophila* systems have made fundamental contributions to understanding *Wolbachia* population biology (e.g., Turelli and Hoffmann 1995; Turelli *et al*. 2022) and the mechanisms of CI (e.g., LePage *et al*. 2017), in addition to modes of *Wolbachia* acquisition (Conner *et al*. 2017; Cooper *et al*. 2019). The second *Wolbachia* found in a *Drosophila* species, *w*Mel, was described in Australian *D. melanogaster* (Hoffmann 1988). Both *w*Ri and *w*Mel rapidly spread worldwide within these invasive hosts (Turelli and Hoffmann 1991; Richardson *et al*. 2012; Kriesner *et al*. 2013, 2016). *w*Ri, which causes strong CI in its native host, now occurs at relatively stable frequencies (∼93%) in almost all characterized *D. simulans* populations (Kriesner *et al*. 2013), while the frequencies of *w*Mel, which causes relatively weak CI in its native host, vary significantly among *D. melanogaster* populations, largely because of variation in the fidelity of maternal transmission (Kriesner *et al*. 2016; Hague *et al*. 2022, 2024). When experimentally transferred from *D. melanogaster* to *Aedes aegypti*, hosts with a most recent common ancestor (MRCA) about 250 MYA (Wiegmann *et al*. 2011), pathogen-blocking *w*Mel causes very strong CI (but see Ross *et al*. 2017). CI facilitates the establishment of *w*Mel in natural *Ae. aegypti* populations and successful biocontrol of dengue (Walker *et al*. 2011; Utarini *et al*. 2021; Lenharo 2023; Velez *et al*. 2023)

Here we focus on determining the timescale of divergence and evolution of *Wolbachia* closely related to *w*Mel (Martinez *et al*. 2021) and observed in holometabolous host species diverged about 350 MY. We present a new temporal calibration of *Wolbachia* divergence using cladogenically inherited *Wolbachia*. Changes of these *Wolbachia* over time scales on the order of 10^6^ years may more accurately represent divergence rates for closely related *Wolbachia* (having diverged on the order of 10^5^ years or less) among distantly related hosts than the mutation-based calibration of Richardson *et al*. (2012). Using our new calibration, we demonstrate that variants closely related to *w*Mel have naturally colonized dipteran and hymenopteran hosts over about the last 370 KY. Many of these *Wolbachia* cause CI and their host invasions are accompanied by even faster turnover of *cifs* among *Wolbachia* genomes. We confirm that *cif* movement occurs with and without the *Wovirus* prophages that contain them, with closely related *cifs* observed in distantly related phages and distantly related *cifs* observed in closely related phages. We also quantify patterns of selection that indicate preservation of CI rescue function and particular Cif protein domains. Our results contribute to a broader understanding of *Wolbachia* and *cif* evolution, while identifying novel *w*Mel-like variants that may serve as candidates for future *Wolbachia* applications.

## Results and discussion

### New time calibration for *Wolbachia* divergence

We present a new conservative calibration for *Wolbachia* chronograms based on examples of *Wolbachia* co-divergence with their hosts. As noted above, the *Nasonia* data of Raychoudhury *et al*. (2009) produced the first example of time-calibrated *Wolbachia*-host co-divergence. The Gerth and Bleidorn (2017) analyses of nuclear and *Wolbachia* genomes across the bee clade (*Nomada ferruginata*, (*N. panzer*, (*N. flava*, *N. leucophthalma*))) provide additional examples. As explained in our Materials and Methods, their data seem most consistent with *Wolbachia* entering the common ancestor of these four species, then diverging between *N*. *ferruginata* and its three-species sister clade. Our analyses below suggest that these latter three species experienced horizontal *Wolbachia* transmission (Conner *et al*. 2017, Table 2; Meany *et al*. 2019, p. 1288). Comparing the *Wolbachia* in *Nomada ferruginata* with those in the sister clade (*N. panzer*, (*N. flava*, *N. leucophthalma*)) suggests a slower rate of *Wolbachia* divergence than the *Nasonia* data, roughly 5.6×10^−10^ [with 95% credible range of (0.40–1.16)×10^−9^] versus 2.2×10^−9^ per year for third sites. To these examples, we add a new calibration based on plausible cladogenic *Wolbachia* transmission between two species in the *Drosophila montium* species group, *Drosophila bicornuta* and *D. barbarae* (Conner *et al*. 2021). This *Drosophila* example implies a substitution rate per year at third sites of 2.2×10^−9^ [with 95% credible range of (1.9–2.6)×10^−9^] very similar to that estimated for *Nasonia*. We use Bayesian analyses to estimate *Wolbachia* chronograms by applying alternative prior distributions based on averaging these examples of codivergence of hosts and *Wolbachia*. The average *Wolbachia* divergence rate for all four of our priors is 1.65×10^−9^ per third site per year, about four times slower than the mutation-based prior from Richardson *et al*. (2012). This longer time-scale does not alter our conclusions here or those of Turelli *et al*. (2018) about “rapid” movement of *Wolbachia* among hosts and the even faster evolution of these closely related *Wolbachia* across those hosts. We expect that the Richardson *et al*. (2012) mutation-rate calibration underestimates divergence times for closely related *Wolbachia* among distantly related hosts, whereas our new calibration, based on co-divergence, may slightly overestimate those times. Our central conclusions are robust to these uncertainties.

**Table 2.** Substitution rate estimates used to calibrate *Wolbachia* chronograms.

| Method | Substitutions/site/year |  | Data | Time calibration/validation |
| --- | --- | --- | --- | --- |
|  | third sites | synonymous sites |  |  |
| “Universal” estimate of the average substitution rate for coding DNA in bacteria applied to *Wolbachia* *ftsZ*<sup>1</sup> | NA | 7–8×10<sup>−9</sup> (ref. 2) | *ftsZ* differences among *Wolbachia*<sup>1</sup> | Divergence-time estimates based on 21 kb of coding DNA from bacteria *Salmonella typhimurium* and *Escherichia coli*<sup>2</sup> supplemented with data indicating similar synonymous-site substitution rates for *ftsZ* in *Wolbachia*<sup>1</sup>. |
| Updated universal synonymous-site substitution rate<sup>3</sup> for bacteria cross-validated with a host nuclear clock<sup>4</sup> for synonymous sites in cladogenically transmitted *Wolbachia* (*wNlonB1* vs. *wNgirB*) in *Nasonia longicornis* and *N. giraulti* | 2.2×10<sup>−9</sup> | 3.3×10<sup>−9</sup> | Fragments of *Wolbachia* and host nuclear loci | Concordance of divergence times estimated from molecular clocks applied to *Wolbachia* and host nuclear DNA<sup>4</sup>. Our substitution-rate estimates assume host and *Wolbachia* diverged 0.46 MYA, averaging the bacterial and eukaryotic clock-based estimates from ref. 4. |
| Divergence of mitochondrial vs. *Wolbachia* genomes among *Drosophila melanogaster* isofemale lines<sup>5</sup> | 6.87×10<sup>−9</sup> | NA | mtDNA and *Wolbachia* genomes | Compare sequence differences among isofemale lines for mtDNA vs. *Wolbachia*. Calibrate substitution rates using the per generation mtDNA mutation rate as a prior for “short term” mtDNA and *Wolbachia* evolution<sup>5</sup>. |
| Cladogenic *Wolbachia* in *Nomada* bees (*N. ferruginata* vs. three-species ingroup – see Table S1) (95% credible intervals based on credible range of divergence times<sup>6</sup>) | 0.56×10<sup>−9</sup>,<br>(0.40–1.16×10<sup>−9</sup>) | 0.91×10<sup>−9</sup><br>(0.65–1.85×10<sup>−9</sup>) | Draft *Wolbachia* and host nuclear genomes | Host divergence-time estimates and 95% credible interval, 2.42 (1.16–3.37) MY, obtained from Fig. 3 of ref. 6, based on refs. 7 and 8. We average the three estimates supported by Table S1. |
| Cladogenic *Wolbachia* in *Drosophila bicornuta* vs. *D. barbara*<sup>9</sup> (95% credible intervals based on credible range of divergence times) | 2.2×10<sup>−9</sup><br>(1.9–2.6×10<sup>−9</sup>) | 3.4×10<sup>−9</sup><br>(2.9–4.0×10<sup>−9</sup>) | Draft *Wolbachia* and host nuclear genomes | Host divergence-time estimates and 95% credible interval, 5.5 (4.65–6.35) MY, from Fig. 1 of ref. 10, |
| **Average estimate from three plausible examples of cladogenic *Wolbachia* transmission** | **1.65×10<sup>−9</sup>** |  |  | Equal weight to three examples: *Nasonia*, *Nomada*, *Drosophila* |

**References:** 1. Werren *et al.* 1995. 2. Ochman and Wilson 1987. 3. Ochman *et al.* 1999. 4. Raychoudhury *et al.* 2009. 5. Richardson *et al.* 2012. 6. Gerth *et al.* 2013. 7. Gerth and Bleidorn 2017. 8. Cardinal *et al.* 2018. 9. This paper. 10. Suvorov *et al.* 2022.

### Rapid spread of *w*Mel-like *Wolbachia* across hosts diverged 350 MYA

To illustrate the timescale of *Wolbachia* movement across host species, we use our new calibration to focus on *Wolbachia* closely related to *w*Mel. Our analyses below also focus on the timescale of *cif* movement and molecular evolution among *Wolbachia* genomes. We use “*w*Mel-like” to designate the clade examined here (Hague *et al*. 2020a), comparing the results to “*w*Ri-like” *Wolbachia* (Turelli *et al*. 2018). We focus on the *Wolbachia* genomes available before Vancaster and Blaxter (2023). Although the clade boundaries are arbitrary, our central conclusions concerning rapid movement of closely related *Wolbachia* among distantly related hosts—and rapid turnover of *Wovirus* and *cifs* within those *Wolbachia* genomes—rest on robustly estimated chronograms and do not require complete sampling of the *Wolbachia* clades we analyze (Supplemental Discussion). When pervasive recombination and horizontal exchange were initially described among diverse *Wolbachia* (Jiggins *et al*. 2001; Baldo *et al*. 2006), it was conjectured that recombination precluded reliable bifurcating phylogenies and chronograms for these mosaic genomes. However, subsequent analyses (cf. Wang *et al*. 2020) show that despite rapid and frequent movements of CI-determining loci, phages, and other genetic elements, large portions of the *w*Mel-like and *w*Ri-like genomes show no significant evidence of recombination (Supplementary Discussion). We use these apparently quasi-clonal genomic regions for our phylogenetic and divergence-time analyses and show that previous analyses (Turelli *et al*. 2018; Meany *et al*. 2019) which did not explicitly control for recombination are robust (Fig. S1; Supplementary Discussion). Note that our analyses do not contradict previous evidence of extensive recombination involving relatively distantly related *Wolbachia*. Rather our analyses of closely related *Wolbachia* show no detectable recombination, as might be expected with relatively rare genetic exchange between nearly identical genomes. These observations are analogous to those concerning horizontal transmission of *Wolbachia*: horizontal transmission is clearly common among distantly related hosts, but it seems quite rare within individual host species (e.g., Richardson *et al*. 2012; Cooper *et al*. 2019).

We consider 20 *w*Mel-like *Wolbachia* whose host species include dipteran and hymenopteran hosts (MRCA: 350 MYA) (Fig. 1A–C) (Wang *et al*. 2016). Using all single-copy genes of equal length with little evidence of past recombination (Supplementary Discussion), the bulk of these *Wolbachia* genomes diverged only about 500 thousand years ago (500 KYA) (95% credible interval: 263 KYA–1.2 MYA) (Fig. 1A). Fig. 1B shows an approximate chronogram for the most diverged hosts: a wasp, *Diachasma alloeum*; a stalk-eyed fly, *Sphyracephala brevicornis*; and a representative drosophilid, *Drosophila teissieri*. The divergence time of the insect orders Diptera, which includes the families Drosophilidae and Diopoidea (stalk-eyed flies) and Hymenoptera is comparable to the crown age of all extant tetrapods, ∼373 million years (MY) (Simões and Pierce 2021). In contrast, the *w*Mel-like *Wolbachia* in these most-diverged hosts, denoted *w*Dal, *w*Sbr and *w*Tei, respectively, diverged about 370 KYA (95% CI: 187–824 KYA) (Fig. 1A node with gray circle). We also report *w*Mel-like *Wolbachia* in 18 drosophilids (Fig. 1C), including a variant, *w*Zts, in tropical *Zaprionus tscasi* that is now the closest known relative of *w*Mel in *D. melanogaster*. *Zaprionus* flies are members of the *Drosophila* subgenus that diverged from the *Sophophora* subgenus about 47 MYA (95% CI: 43.9–49.9 MYA), highlighting *w*Mel proliferation across species that span the entire paraphyletic genus also named *Drosophila* (Suvorov *et al*. 2022). The drosophilid host range and rapidity of movement for *w*Mel-like variants are similar to estimates for *w*Ri-like variants (Turelli *et al*. 2018), suggesting that this “life history” may characterize many common *Wolbachia*. In our Supplementary Discussion, we elaborate additional inferences that can be made when more than one *Wolbachia* sequence from these hosts and others become available and provide a correction of prior claims concerning the *Wolbachia* in *D. suzukii* and *D. subpulchrella* based on species misidentification (Fig. S2, Supplementary Discussion). The complete set of hosts and their *w*Mel-like *Wolbachia* is provided in Table S1.

**Figure 1.**
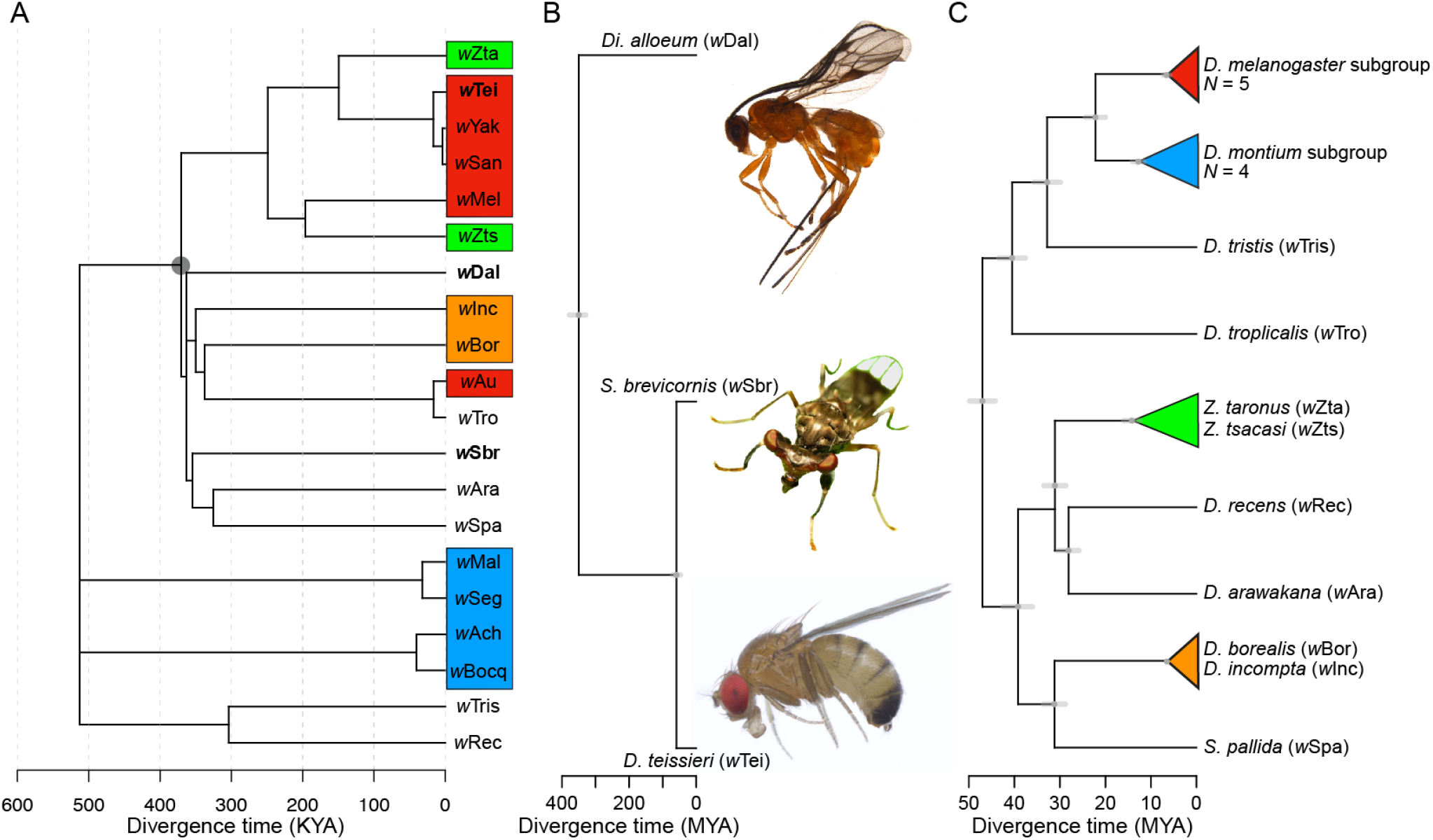
*w*Mel-like *Wolbachia* that diverged approximately 370 KYA occupy insects that diverged about 350 MYA. **(A)** An absolute chronogram with the *Wolbachia* associated with the most distantly related hosts in bold. The colored *Wolbachia* labels match the host clades from Panel C. The crown age is 512 KY, with 95% credible interval of 263 K to 1.2 MY. **(B)** An approximate chronogram for the most distantly related host clades containing *w*Mel-like *Wolbachia*: Hymenoptera (*Diachasma alloeum*) and within Diptera, Diopsidea (*Spyracephala brevicornis*) and Drosophilidae (*D. teissieri* presented as a representative drosophilid). Diptera and Hymenoptera diverged about 350 MYA (378–329 MYA, Devonian–Carboniferous) and span Holometabola (Misof et al. 2014; Johnson et al. 2018). Drosophilidae and the Diopsoidea superfamily containing Diopsidea stalk-eyed flies diverged about 59 MYA based on the crown age of the Drosophilidae (47 MY) (Suvorov *et al*. 2022), and the crown age of Schizophora (70 MY) (Wiegmann *et al*. 2011). The *w*Mel-like *Wolbachia* in these most-diverged hosts, denoted *w*Dal (in *D. alloeum*), *w*Sbr (in *S. brevicornis*) and *w*Tei (in *D. teissieri*) in bold in Fig. 1A diverged about 370 KYA (95% CI: 187–824 KYA; Fig. 1A node denoted with gray circle) **(C)** A chronogram for drosophilid hosts with node ages and approximate confidence intervals estimated from the fossil-calibrated chronogram of Suvorov *et al*. (2022). Images taken by Centre for Biodiversity Genomics (*D. alloeum*), Katja Schulz (*S. brevicornis*) and Tim Wheeler (*D. teissieri*).

### Rapid introgressive transfer of *Wolbachia* between some closely related species

Many obligate mutualistic endosymbionts like the *Wolbachia* in filarial nematodes (Comandatore *et al*. 2013) and *Buchnera* in aphids (Baumann *et al*. 1995) are acquired cladogenically. In contrast, all but one of these *w*Mel-like *Wolbachia* must have been acquired through introgression or non-sexual horizontal transmission. Introgression is plausible only between the most closely related drosophilid species in our study (indicated by the colored triangles in Fig. 1C). Joint analysis of mtDNA and *Wolbachia* sequences implies that the three-species *yakuba* clade (*D. teissieri*, (*D. yakuba*, *D. santomea*)) first acquired *Wolbachia* by horizontal transmission from an unknown host, then transferred it within the clade through hybridization and introgression (Cooper *et al*. 2019). *D. yakuba* hybridizes with its endemic sister species *D. santomea* on the island of São Tomé (Lachaise *et al*. 2000; Comeault *et al*. 2016; Cooper *et al*. 2017), and with *D. teissieri* on the edges of forests on the nearby island of Bioko (Cooper *et al*. 2018). *Wolbachia* and mtDNA chronograms are generally concordant for these three hosts and indicate more recent common ancestors for these maternally inherited factors than for the bulk of the host nuclear genomes (Cooper *et al*. 2019). *Z. taronus* occurs on São Tomé, particularly co-occurring with *D. santomea* at high altitudes; but it diverged from the *D. yakuba* triad about 47 MYA (Fig. 1C), making introgression impossible. Yet, its *Wolbachia* (*w*Zta) diverged from the *w*Yak-clade *Wolbachia* only about 54–353 KY. These data illustrate horizontal *Wolbachia* transfer between distantly related species with overlapping ranges and habitats.

Other possible examples of introgressive *Wolbachia* acquisition involve two species pairs in the *D. montium* subgroup, (*D. seguyi*, *D. malagassya*) and (*D. bocqueti*, *D.* sp. aff. *chauvacae*), whose *Wolbachia* diverged on the order of 32 KY (11–87 KY) and 40 KY (14–106 KY), respectively. Both host pairs appear as sister species (Conner *et al*. 2021), so introgressive *Wolbachia* transfer is plausible. However, the mtDNA third-position coding sites differ by 0.53% and 1.15% respectively, corresponding to divergence times on the order of 100 KY or longer (Ho *et al*. 2005). Our estimate of *w*Seg-*w*Mal divergence is inconsistent with introgressive acquisition by *D. seguyi* and *D. malagassya*, while we cannot rule out introgressive acquisition by *D. bocqueti* and *D.* sp. aff. *chauvacae* based on our credible interval of *w*Bocq-*w*Ach divergence. For the more distantly related pairs ((*Z. taronus*, *Z. tsacasi*) and (*D. borealis*, *D. incompta*)), the mtDNA third-site differences of 19.3% and 30% respectively, decisively preclude introgressive *Wolbachia* transfer.

### *w*Mel-like *Wolbachia* hosts are diverse and cytoplasmic incompatibility is common

The hosts of these *w*Mel-like *Wolbachia* are extraordinarily diverse in ecology and geography. They range from cosmopolitan human-associated species (*D. melanogaster*, *D. simulans* and *S. pallida*) to endemics restricted to small oceanic islands (*D. santomea* and *D. arawakana*). The drosophilids include one that breeds and feeds on flowers (*D. incompta*), a mushroom specialist (*D. recens*), and classic generalists (e.g., *D. melanogaster* and *D. simulans*). As expected, hosts with closely related *Wolbachia* co-occur somewhere (or did in the recent past). For instance, *w*Au was previously observed in *D. simulans* in Florida and Ecuador (Turelli and Hoffmann 1995). Thus, before *w*Au was displaced by *w*Ri, *w*Au-carrying *D. simulans* probably co-occurred with *D. tropicalis* found only in Central and South America and Caribbean islands, which harbors *w*Tro, sister to *w*Au in Fig.1C. Although the wasp *Diachasma alloeum* parasitizes the tephritid *Rhagoletis pomonella*, none of *R*. *pomonella*’s several *Wolbachia* seem to be *w*Mel-like (Schuler *et al*. 2011). We focus on the phylogenetic distribution of CI-inducing *Wolbachia* associated with these hosts.

Of the 20 *w*Mel-like and 8 *w*Ri-like *Wolbachia* in our study, all but 11 *w*Mel-like strains have been tested for CI. For 2 of these 11 *w*Mel-like strains (*w*Seg in *D. seguyi* and *w*Bocq in *D. bocqueti*), insect stocks were available for us to test for CI. Putatively incompatible conspecific crosses between females without *Wolbachia* and males with *Wolbachia* (IC) produce lower egg hatch than do conspecific compatible crosses (CC) for both *w*Seg in *D. seguyi* (IC egg hatch = 0.34 ± 0.21 SD, *N* = 14; CC egg hatch = 0.89 ± 0.11 SD, *N* = 18; *P* < 0.001) and *w*Bocq in D. *bocqueti* (IC egg hatch = 0.18 ± 0.24 SD, *N* = 13; CC egg hatch = 0.60 ± 0.30 SD, *N* = 17; *P* = 0.002). This confirms relatively strong CI in two new *w*Mel-like *Wolbachia* systems. In total, 8 *w*Mel-like and 6 *w*Ri-like *Wolbachia* in our study cause CI (Fig. 2A; see Table S1 for references). CI strength varies greatly among these systems and others (Hoffmann 1988; Cooper *et al*. 2017; Shropshire *et al*. 2022), but may also vary within systems (Shropshire *et al*. 2020) as a function of male age (Reynolds and Hoffmann 2002; Shropshire *et al*. 2021b), environmental factors (Clancy and Hoffmann 1998; Ross *et al*. 2017), and host backgrounds (Poinsot *et al*. 1998; Reynolds and Hoffmann 2002; Cooper *et al*. 2017; Wybouw *et al*. 2022). This includes *w*Mel, that tends to express weak CI in *D. melanogaster* (Hoffmann 1988) and strong CI in other hosts (Zabalou *et al*. 2008; Walker *et al*. 2011). *Wolbachia* that carry putatively functional *cifs*, but that do not cause CI in their natural hosts, are candidates for future work focused on the evolution of host suppression of CI (see below).

**Figure 2.**
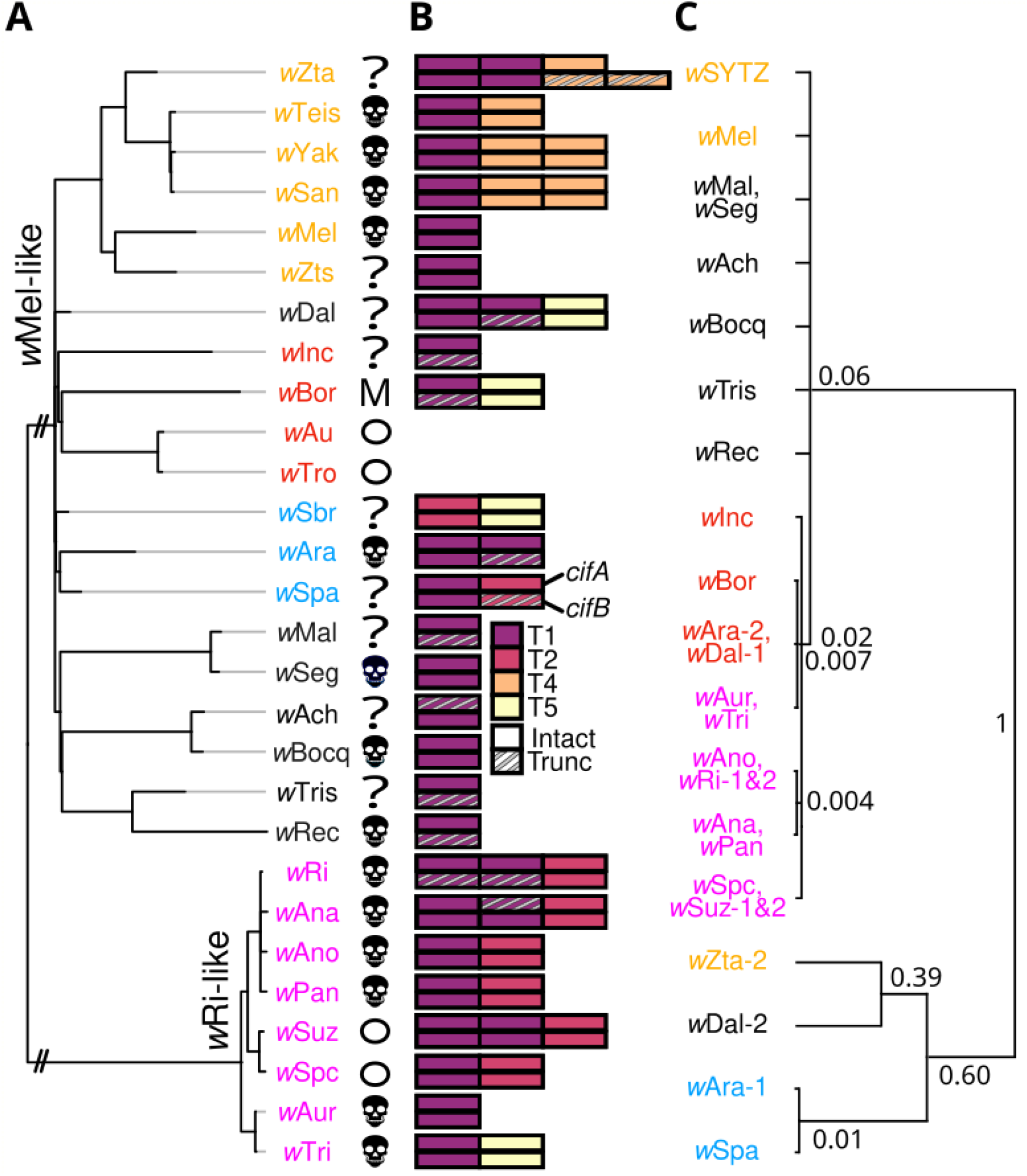
Diverse *cif* operons rapidly turnover among *w*Mel-and *w*Ri-like genomes. **(A)** A phylogram of *w*Mel-like and *w*Ri-like *Wolbachia*, including variants that cause CI (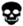), do not cause CI (circles), or have unknown CI status (?). *w*Bor does not cause CI, but it does kill males (M). The *w*Mel-like and *w*Ri-like clades diverged 2–10.4 MYA (see Supplementary Discussion). Branches leading to these clades are shortened (//) and light gray branch extensions are used to improve visualization. **(B)** These closely related *Wolbachia* carry four of five known *cif* operon Types (T1– T5). *cifA* (top) and *cifB* (bottom) schematics are presented with operon copies adjacent to one another. **(C)** A relative chronogram for *cifA_[T1]_* with node labels indicating relative ages, scaled to 1 for the most diverged. Identical sequences are collapsed into a single tip, and nodes with posterior probability < 0.95 are collapsed into polytomies. Strain labels are colored to highlight *Wolbachia*-*cifA* discordance.

### *cif* genes and proteins are highly diverse among *w*Mel-like *Wolbachia*

CI is caused by two-gene *cif* operons, with paternal expression of *cifB* (and occasionally *cifA*) killing embryos unless a complementary *cifA* copy is expressed maternally (LePage *et al*. 2017; Beckmann *et al*. 2017; Shropshire *et al*. 2018, 2021a; Shropshire and Bordenstein 2019; Xiao *et al*. 2021; Adams *et al*. 2021; Sun *et al*. 2022). *cifs* are generally associated with *Wovirus* bacteriophages that are themselves subdivided into four groups typed using serine recombinase (sr) alleles (sr1WO–sr4WO), with three containing *cifs* (sr1WO-sr3WO) (Bordenstein and Bordenstein 2022). *Wolbachia*-encoded *cifs* span five described clades called Types (*cif*_[T1]_–*cif*_[T5]_) (Martinez *et al*. 2021), and *Wolbachia* genomes often contain multiple *cif* operons (Fig. 2B) (Bonneau *et al*. 2018; Martinez *et al*. 2021). Excluding non-CI-inducing *w*Mel-like *w*Au and *w*Tro (Turelli and Hoffmann 1995; Hoffmann *et al*. 1996; Martinez *et al*. 2015), the *Wolbachia* in our analyses encode between one and three *cif* operons from four of the five described *cif* types: all genomes except *w*Sbr contain a *cif*_[T1]_ operon, eight contain a *cif*_[T2]_ operon, and *cif*_[T4]_ and *cif*_[T5]_ operons each appear in four *Wolbachia* (Fig. 2B). Even within *cif* types, there is significant variation in protein sequence, length, and domain composition (Fig. S3A-C; Supplementary Discussion). An exceptional example is CifB*_w_*_Zta[T1-2]_ (i.e., the second copy of CifB_[T1]_ found in *w*Zta). It is 357 amino acids longer than the next largest CifB_[T1]_ and shares only 51% to 39% sequence identity with other CifB_[T1]_ proteins (Table S2). Despite its considerable divergence, *w*Zta’s CifB is more similar to other CifB_[T1]_ proteins in our dataset than to any other putatively functional CifB types, and retains a pair of PD-(D/E)XK nuclease domains ubiquitous among CifB proteins (Kaur *et al*. 2024).

### *cif* operons turn over rapidly among closely related *Wolbachia*

*Wolbachia* and *cif* phylogenies are often discordant (LePage *et al*. 2017; Cooper *et al*. 2019; Martinez *et al*. 2021), but the timing of *cif* movement among *Wolbachia* genomes is unresolved. Using our new calibration and comparison of *cif* operons observed in closely and distantly related *Wolbachia* genomes, we document rapid turnover of multiple *cif* types. Several *w*Mel-like clades provide clear evidence of *cif* turnover. In the *w*SYTZ clade (*w*SYT plus the *w*Zta outgroup), *w*SYT genomes contain only one *cif*_[T1]_ operon but *w*Zta contains two, indicating a gain or loss in the last 54–353 KY (see below, Fig. 2). Sequence divergence between *w*Zta’s two *cif*_[T1]_ operons implies that acquisition of the second operon did not involve duplication (Fig. 2C). Turnover is not restricted to particular *cif* types, as exemplified by the *w*SYTZ clade acquiring *cif*_[T4]_ operons after diverging from (*w*Mel, *w*Zts) 117–569 KYA (Supplementary Discussion). *w*SYT *Wolbachia* also differ in *cif*_[T4]_ copy number, with *w*Yak and *w*San acquiring a second copy since diverging from *w*Tei (MRCA: 4.8**–**44 KYA) (Baião *et al*. 2021). In the ((*w*Ara, *w*Spa), *w*Sbr) clade (MRCA: 170–791 KYA), we observe turnover of *cif*_[T1]_, *cif*_[T2]_ and *cif*_[T5]_ operons. *w*Ara contains two *cif*_[T1]_ operons, its sister *w*Spa contains a *cif*_[T1]_ operon and a *cif*_[T2]_ operon, and outgroup *w*Sbr contains only *cif*_[T2]_ and *cif*_[T5]_ operons. In the (*w*Inc, (*w*Bor, (*w*Au, *w*Tro))) clade (MRCA: 179–790 KYA), *w*Inc and *w*Bor contain a *cif*_[T1]_ operon, *w*Bor contains a *cif*_[T5]_ operon, and *w*Au and *w*Tro do not contain any operons. *cif* turnover is not restricted to *w*Mel-like *Wolbachia* as exemplified by the *w*Ri-like *w*Tri genome containing a *cif*_[T5]_ operon that is absent in the very closely related *w*Aur sister variant (no observed differences across the 525 genes and 506,307 bp used to produce the *w*Ri-like phylogram in Turelli *et al*. (2018).

We further illustrate rapid *cif* turnover by focusing on homologs of *cifA*_[T1]_, which is the most common *cif* type in our dataset (Fig. 2C, Supplementary Discussion). We observe two distantly related clades of *cifA*_[T1]_ alleles. The first includes 25 alleles observed across 15 *w*Mel-like *Wolbachia* and 8 *w*Ri-like *Wolbachia. w*Ri and *w*Suz each carry two closely related *cifA*_[T1]_ alleles, likely originating via duplication. *cifA*_[T1]_ alleles observed in *w*Mel-like *w*Dal, *w*Ara, *w*Bor and *w*Inc genomes are most closely related to *cifA*_[T1]_ alleles observed in *w*Ri-like *Wolbachia* genomes and share particularly high identity with *cifA*_[T1]_ alleles observed in *w*Ri-like *w*Aur and *w*Tri genomes (99.8–99.2% aa identity). The second *cifA*_[T1]_ clade includes additional *cifA*_[T1]_ copies in *w*Mel-like *w*Zta, *w*Dal, and *w*Ara genomes that are more closely related to each other than each is to the second *cifA*_[T1]_ copy they carry. This clade also contains a *cifA*_[T1]_ allele observed in *w*Spa (*w*Spa, *w*Ara). While we cannot resolve the complete history of *cif* movement and evolution from these data, we conclude that *cif* turnover is rapid, occurring within and between *w*Mel-like and *w*Ri-like clades on the order of 10^4^-10^5^ years.

### *Wovirus* turnover does not fully explain *cif* movement

Two additional aspects of our data on *cif* transfer are worth emphasizing. First, the *Wovirus* bacteriophages that contain *cifs* also rapidly turn over among *Wolbachia* genomes (Supplementary Discussion). An exceptional case involves the *w*Bocq genome that contains an sr3WO *Wovirus* that is absent from the genome of sister *w*Ach. This implies *Wovirus* gain or loss in the last 14–106 KY. Second, while *cifs* clearly transfer along with the *Wovirus* that carry them, *cifs* also move among divergent *Wovirus* classes (i.e., phage-independent *cif* turnover) (Cooper *et al*. 2019; Baião *et al*. 2021). This interchange is documented in two ways: closely related *cifs* are found in distantly related phages and distantly related *cifs* are found in closely related phages (Fig. 3, and Supplementary Discussion). We demonstrate this by comparing phylograms of the serine recombinase genes (sr) of sr3WO *Wovirus* (sr3) to phylograms of *cifA*_[T1]_ alleles associated with them. The ten sr3WO *Wovirus* in *w*Ri-like *Wolbachia* have identical sr3 alleles and are closely related to sr3 alleles found in several *w*Mel-like *Wolbachia*. These sr3 alleles are more distantly related to several other sr3 alleles that include a copy found in *w*Mel from *D. melanogaster*. In contrast, almost all *cifA*_[T1]_ alleles associated with *w*Mel-like sr3WO *Wovirus* are very closely related to *cifA*_[T1]_ alleles associated with *w*Ri-like sr3WO *Wovirus* (Fig. 3). This generalizes phage-independent *cif* transfer among divergent *Wolbachia* that was first documented for Type IV loci in *w*Yak by Cooper *et al*. (2019) and later misinterpreted by Baião et al. (Baião *et al*. 2021) (Supplementary Discussion). Higher quality assemblies for known donor and recipient *Wolbachia* will be essential for establishing the relative role of insertion sequence (IS) elements (Cooper *et al*. 2019) and other factors like plasmids in this transfer. We hypothesize a critical role for IS elements in the phage-independent *cif* transfer we document, as supported by IS elements flanking the majority of *cifs* in our analyses (Table S3).

**Figure 3.**
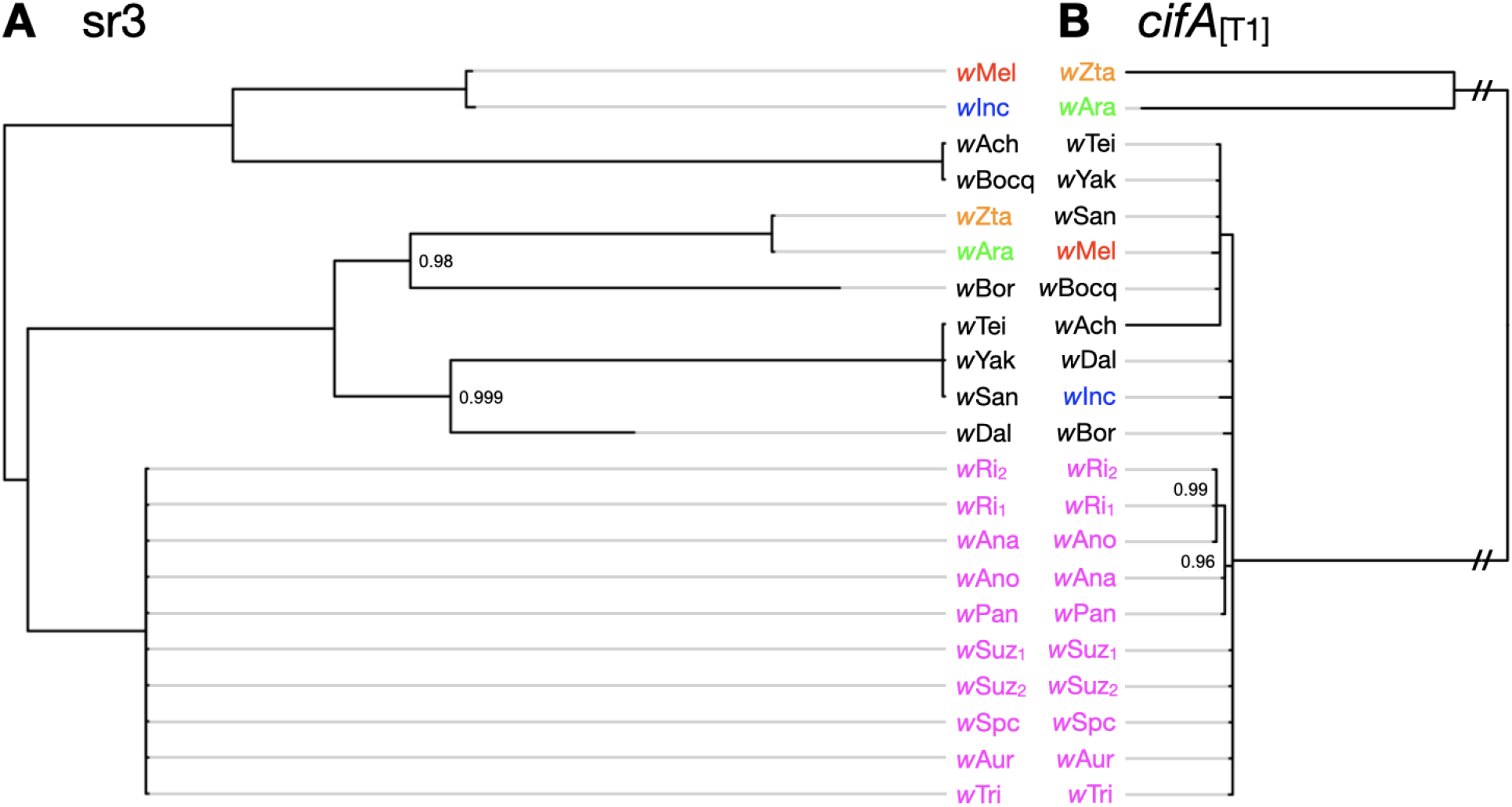
Discordant phylograms for sr3 alleles of sr3WO and linked *cifA_[T1]_* alleles demonstrate phage-independent cif turnover. **(A)** Phylogram for sr alleles facing **(B)** phylogram for *cifA_[T1]_* alleles linked to these sr3 alleles. Branches leading to the two sets of closely related *cifA_[T1]_* alleles are shortened (//) to improve visualization. *w*Ri-like strains are shown in magenta, and focal *w*Mel-like strains are colored to highlight sr3-*cifA_[T1]_* discordance. Subscripts represent different sr3 copies within the same *Wolbachia* genome (see Table S3) in cases where multiple sr3 alleles can be associated with specific *cifA_[T1]_* copies. Subscripts presented for *cifA_[T1]_* alleles denote associated sr alleles. Light gray branch extensions are provided to simplify sr3-*cifA_[T1]_* comparisons. Nodes with posterior probability < 0.95 are collapsed into polytomies. Posterior support values appear only at nodes with support less than 1.

### Selection acts to preserve *cifA* and nuclease domains within *cifB*

What is the fate of these *cifs*? Theory predicts that once *Wolbachia* infections are established in a host species, natural selection does not act to maintain CI but does act to maintain resistance to CI (Turelli 1994; Haygood and Turelli 2009). Consistent with these predictions and prior observations (Meany *et al*. 2019; Martinez *et al*. 2021), Fig. 2B shows that putative pseudogenization (i.e., truncation) is more common for *cifB* than for *cifA* (see also Fig. S4A,B). Still, as noted by Beckmann *et al*. (2021), CI is incredibly common despite weak selection on the phenotype (Turelli *et al*. 2022). This paradoxical prevalence of CI across *Wolbachia* lineages can be explained in part by clade selection in which CI-causing *Wolbachia* lineages are more likely to be transmitted to new host species because they typically have higher frequencies within host species and persist longer than do non-CI causing *Wolbachia* (Turelli *et al*. 2022). However, CifB also contributes to alternative functions that include regulation of *Wolbachia* abundance in host tissues through interactions with host autophagy (Deehan *et al*. 2021). This suggests that non-CI pleiotropic effects could plausibly contribute to persistence of particular *cifs*.

To assess patterns of selection across Cifs, we calculated the ratio of non-synonymous (*d_N_*) to synonymous substitutions (*d_S_*) for each Cif protein, using a 3-dimensional spherical sliding window across the length of AlphaFold-derived Cif structures (Fig. 4A, Fig. S3, Fig. S4C,D, Movie S1). Fig. 4B shows that CifA_[T1]_ proteins are more similar to one another in terms of both sequence identity and structural similarity than CifB_[T1]_ proteins from the same pairs. As predicted, CifA of Types 1 and 2 had lower *d_N_*/*d_S_* ratios than did CifB of the same type (Fig. 4, S3D,E) (Supplementary Discussion), consistent with purifying selection maintaining CifA. Putatively pseudogenized CifA and CifB proteins have higher *d_N_*/*d_S_* ratios than do intact proteins (e.g., CifA *d_N_*/*d_S_ ∼* 1; Fig. 4E,F), further supporting the presumption that in-frame stop codons interfere with Cif function (Supplementary Discussion). In contrast, Types 3 and 4 CifA and CifB have comparable *d_N_*/*d_S_* ratios within each type, which could plausibly stem from pleiotropy or loss-of-function (Fig. S3D,E). CifA_[T1]_ and CifB_[T1]_ binding residues and CifB_[T1]_’s Deubiquitinase domain have *d_N_*/*d_S_* ratios comparable to non-domain associated residues. However, the two CifB_[T1]_ nuclease domains both have lower *d_N_*/*d_S_* ratios than do other residues (Fig. 4C,D,G,H). Indeed, across Cif Types, CifB’s first nuclease domain has lower *d_N_*/*d_S_* ratios than non-domain associated residues (Fig. S3F-H). While CifB’s nuclease activity may not always contribute to observed CI expression (Kaur *et al*. 2024), selection may still act to maintain other nuclease-associated features (e.g., DNA binding) and contribute to CifB’s association with chromatin restructuring (Supplementary Discussion) (Kaur *et al*. 2022; Terretaz *et al*. 2023). Interestingly, our data reveal that sites with *d_N_*/*d_S_* ratios above 1, consistent with positive selection, are primarily located within regions of the protein that are not domain-associated (Fig. 4D). Many of these sites appear at the surface, aligning with the expectation that surface residues are key to host-microbe interactions, potentially facilitating interactions between Cif and host proteins. Thus, while theory predicts that selection does not act on the CI phenotype (Turelli 1994), selection on alternative CifB functions may plausibly delay mutational disruption of *cifB* (Beckmann *et al*. 2021).

**Figure 4.**
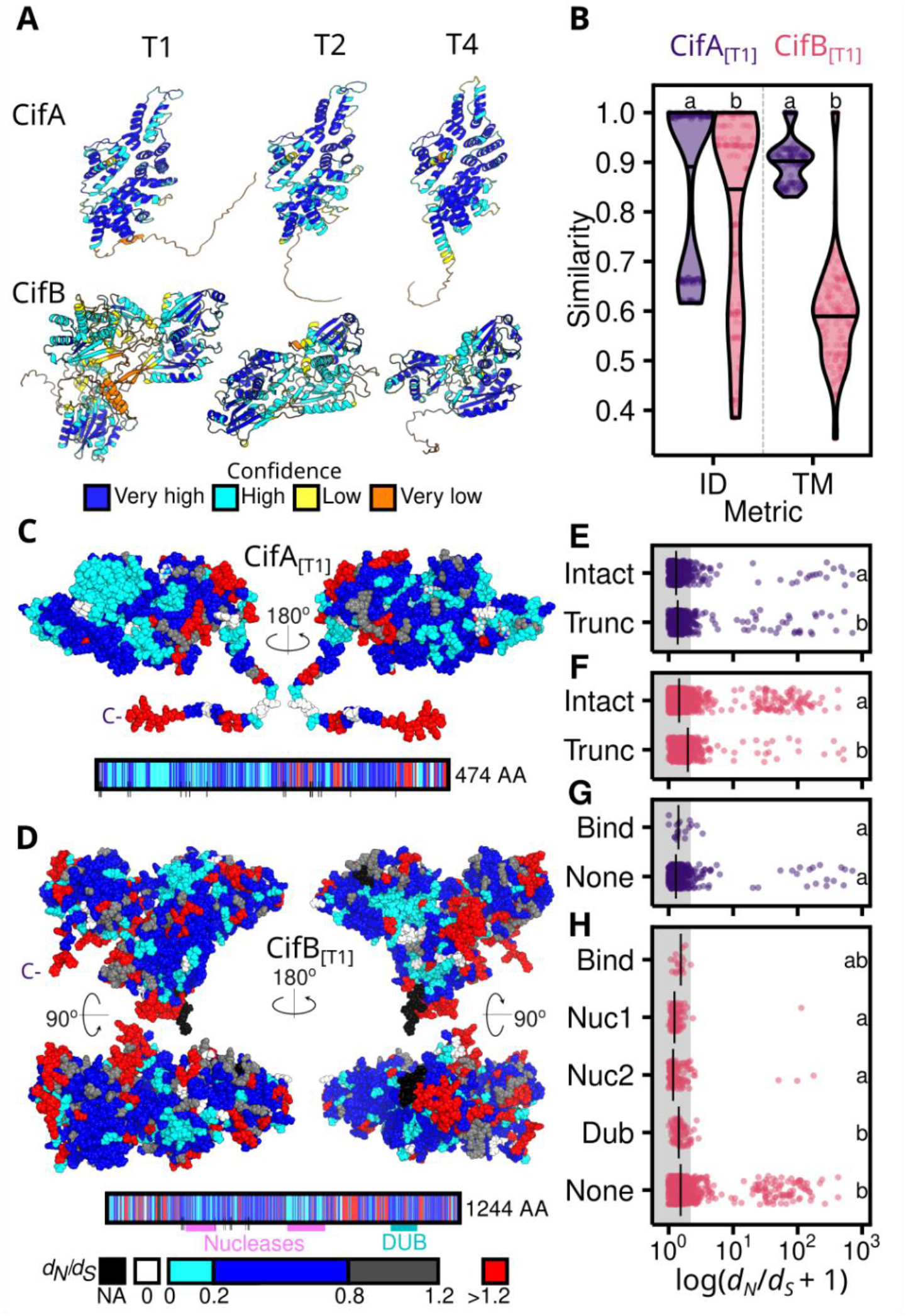
Cifs are highly variable and CifA tends to have lower values of *d_N_/d_S_* than CifB. **(A)** Representative Cif*_w_*_Mel[T1]_, Cif*_w_*_Ri[T2]_, and Cif*_w_*_Yak[T4]_ AlphaFold structures colored by confidence (pLDDT) per residue. **(B)** Putatively intact CifA_[T1]_ proteins are more similar than CifB_[T1]_ from the same pairs (*N* = 18) in terms of both sequence identity (ID) and structural similarity (TM). *d_N_/d_S_* for **(C)** CifA_[T1]_ and **(D)** CifB_[T1]_ displayed on Cif*_w_*_Mel[T1]_ AlphaFold structures and linear schematics. Median *d_N_/d_S_* per residue is calculated using a 10 Å spherical sliding window in pairwise comparisons of Cif*_w_*_Mel[T1]_ to other Cif_[T1]_ proteins (CifA: *N* = 9; CifB: *N* = 4). Colored boxes and black lines below schematics indicate domains and CifA-CifB binding sites, respectively. *d_N_/d_S_* is relatively lower for intact than truncated **(E)** CifA_[T1]_ and **(F)** CifB_[T1]_. **(G)** *d_N_/d_S_* for CifA_[T1]_ binding sites tends to not differ from *d_N_/d_S_* for other residues. **(H)** *d_N_/d_S_* tends to be lower for CifB_[T1]_ nuclease domains than for other residues. Shared letters within plots b and e-h represent statistically similar groups determined by a Mann-Whitney U test (2 groups) or a Kruskal-Wallis and Dunn’s multiple comparison test (>2 groups). *P*-values are presented in Table S4.

### Conclusions

Our findings confirm that non-sexual horizontal *Wolbachia* acquisition—and introgressive transfer between close relatives—commonly occur on the order of 10^4^ to 10^5^ years. These conclusions are robust to uncertainties about *Wolbachia* divergence times. Among recently diverged *Wolbachia*, distantly related *cif* operons can be gained and/or lost even faster, both with and without the *Wovirus* that contain them. The commonness of non-sexually acquired *Wolbachia* and *cif* transfer among *Wolbachia* genomes indicates that opportunities for horizontal transfer must occur often. Because many insects carry vertically transmitted *Wolbachia* (Weinert *et al*. 2015), we expect that ephemeral somatic infections are common (cf. Towett-Kirui *et al*. 2021). Future analyses should focus on understanding the ecology of non-sexual *Wolbachia* transfer and the cellular-genetic basis of successful non-sexual *Wolbachia* establishment (or not) in new hosts, as well as the mechanisms of *cif* transfer with and without *Wovirus* (Cooper *et al*. 2019; Baião *et al*. 2021).

CI-causing *Wolbachia* provide a practical mechanism for mitigating human diseases. While *w*Mel introductions into *Ae. aegypti* have been very effective in locations where they have spread to high, stable frequencies (Hoffmann *et al*. 2011; Utarini *et al*. 2021; Lenharo 2023; Velez *et al*. 2023), alternative *Wolbachia*-based interventions are needed. For example, *w*Mel has been lost in some release locations (Hien *et al*. 2021; Moledo Gesto *et al*. 2021), particularly under extremely hot conditions. Temperature can affect CI strength (Reynolds and Hoffmann 2002; Ross *et al*. 2019) and rates of imperfect *Wolbachia* transmission (Ross *et al*. 2017; Hague *et al*. 2022, 2024); and the temperatures hosts prefer and their overall activities differ when they carry *Wolbachia* (Hague *et al*. 2020b). Identifying strong-CI-causing and virus-blocking *Wolbachia* from the tropics could facilitate *Wolbachia* biocontrol, and *w*Mel-like variants that naturally associate with tropical host species are obvious candidates (Gu *et al*. 2022). Our study expands a comprehensive panel of *w*Mel-like *Wolbachia* and quantifies the timescale of their movement and evolution—these *Wolbachia* exhibit diverse ecology, geography and *cif* profiles. We report that a tropical *Wolbachia* variant (*w*Zts) is now the most closely related known variant to *w*Mel in *D. melanogaster*. While *w*Zts CI has not yet been tested, all six *Z. tscasi* sampled in nature carry *w*Zts (producing 0.61 as the 95% lower bound for *w*Zts frequency), consistent with strong CI. Identification of such variants adds versatility and contributes towards the development of more customized and environment-specific *Wolbachia* applications.

## Materials and methods

### *Wolbachia* assembly and phylogeny estimation

The *Wolbachia* genomes not novel to this study were obtained from the sources listed in Table S1. For the genomes novel to this study or where the source accession is an SRR number (raw reads from NCBI’s SRA database), we trimmed the reads with Sickle v. 1.3 (Joshi and Fass) and assembled them with ABySS v. 2.2.3 (Jackman *et al*. 2017) with Kmer values of 51, 61, …, 91. From these host assemblies, scaffolds with best nucleotide BLAST matches to known *Wolbachia* sequences, with E-values less than 10^−10^, were extracted as the *Wolbachia* assembly. For each host, the best *Wolbachia* assembly (fewest scaffolds and highest N50) was kept. To assess the quality of these draft assemblies, we used BUSCO v. 3.0.0 to search for orthologs of the near-universal, single-copy genes in the BUSCO proteobacteria database. As controls, we performed the same search using the reference genomes for *w*Ri, *w*Au, *w*Mel, *w*Ha and *w*No.

To identify genes for the phylogenetic analyses, all *w*Mel-like and wRi-like *Wolbachia* genomes (see Table S1) were annotated with Prokka 1.14.5 (Seemann 2014), which identifies orthologs to known bacterial genes. To avoid pseudogenes, paralogs and frameshifts, we used only genes present in a single copy in each genome and required that at most one draft genome had gaps in the alignment. Genes were identified as single-copy if they uniquely matched a bacterial reference gene identified by Prokka and aligned with MAFFT v. 7 (Katoh and Standley 2013). We estimated three separate phylogenies. For the set of the 20 *w*Mel-like genomes, 438 genes, a total of 409,848 bp, met these criteria. For the set of 20 *w*Mel-like genomes plus the 8 *w*Ri-likes, 346 genes, a total of 310,977 bp, met these criteria. For the set of 8 *w*Ri-like genomes above and here, we used the same data set used in Turelli *et al*. (2018), which was 525 genes, 506,307 bp. These genes also show little evidence for recombination (see below).

### *Wolbachia* chronograms

To estimate absolute chronograms for the three datasets discussed above—*w*Mel-like clade (Fig. 1A), *w*Ri-like clade (Turelli *et al*. 2018)—and both clades combined—we first estimated a relaxed-clock relative chronogram with RevBayes 1.1.1 (5) with the root age fixed to 1 using the GTR + Γ nucleotide-substitution model, partitioned by codon position, and the same birth-death process prior for tree shape as in Turelli *et al*. (2018). There were not enough substitutions to partition by gene as well as codon position. Each partition had an independent rate multiplier with prior Γ(1,1), as well as stationary frequencies and substitution rates drawn from flat, symmetrical Dirichlet distributions. For each branch, the branch-rate prior was Γ(7,7), normalized to a mean of 1 across all branches. Four independent runs were performed, which always agreed. Nodes with posterior probability less than 0.95 were collapsed into polytomies. In Turelli *et al*. (2018), we converted the relative chronogram constructed as described above into an absolute chronogram using the scaled distribution prior Γ(7,7) × 6.87 × 10^−9^ substitutions per third-position site per year, derived from the mutation-based posterior distribution estimated by Richardson *et al*. (2012). The mean assumes 10 generations per year. Absolute branch lengths were calculated as the relative branch length times the third position rate multiplier estimated from the substitutions per third-position site per year. Here we provide an alternative calibration for the third-position rate based on examples of cladogenic *Wolbachia* transmission. To explore the robustness of our estimates of *Wolbachia* divergence times to the specific probability distributions used as priors, we consider four alternatives discussed below.

Turelli *et al*. (2018) derived an absolute chronogram from their relaxed-clock relative chronogram using a substitution-rate estimate of Γ(7,7)×6.87×10^−9^ substitutions/3rd position site/year. The gamma distribution was used to approximate the variation in the mutation-rate estimate obtained by Richardson *et al*. (2012). Although Γ(7,7) approximated the per-generation variance estimate, Turelli et al. (2018) used it to approximate the per-year variance. Because Turelli *et al*. (2018) assumed 10 generations per year, the variance per year was underestimated in this analysis. Here we use a different approach to approximating the uncertainty in substitutions/third-position site/year. Our new variance estimates depend on estimated variation in the divergence times for the host-*Wolbachia* pairs used to calibrate *Wolbachia* DNA divergence rates (see Tables 1 and 2).

The first three rows of Table 1 summarize data describing the divergence of *Wolbachia*, nuclear loci and mtDNA loci across three plausible examples in which *Wolbachia* codiverged with their hosts. Particularly notable is the relative consistency of the ratios of *Wolbachia* versus nuclear sequence divergence. The next six rows of Table 1 show the comparable data for the *Nomada* studied by Gerth and Bleidorn (2017). When the outgroup species, *N. ferruginata*, and its *Wolbachia* are compared pairwise to the nuclear genomes and *Wolbachia* of the three ingroup species, (*N. panzeri*, *N. flava* and *N. leucophtalma*), the ratios of *Wolbachia* to nuclear divergence are broadly consistent with those estimated from the *Nasonia*, *Drosophila* and *Brugia* examples. In contrast, the pairwise divergences estimated within the ingroup (*N. panzeri*, *N. flava* and *N. leucophtalma*) suggest a relative *Wolbachia* to nuclear divergence rate that is about 5–20 times faster than the other putative examples of cladogenic *Wolbachia* transmission. We suspect that *Wolbachia* may have been horizontally transferred within the three-species *Nomada* ingroup. To produce a more conservative estimate of the time scale of *Wolbachia* horizontal movement and evolution, we use only the three comparisons between *N. ferruginata* and the other three *Nomada* species in our new *Wolbachia* calibration presented in Table 2.

For their *Nasonia* analyses, Raychoudhury *et al*. (2009) used widely applied eukaryotic and bacterial molecular clock calibrations to separately estimate the divergence times for hosts and their *Wolbachia*. We averaged those estimated divergence times to produce a divergence-time estimate of 0.46 million years (MY). That estimate produces an average third-site *Wolbachia* substitution rate of 2.2×10^−9^ substitutions/third-position site/year over the 4486 bp of their *Wolbachia* DNA sequence data (see Table 2). For our *Drosophila* and *Nomada* pairs with plausible cladogenic *Wolbachia* transmission, we have fossil-based estimates of the host divergence times that do not rely on molecular clock approximations. For the *Drosophila* pair, *D. bicornuta* and *D. barbarae*, Suvorov *al et.* (2022, Fig. 1) provides a divergence-time estimate of 2.42 MY, with 95% credible interval of 1.16–3.37 MY. Using 620,685 bp extracted from single-copy *Wolbachia* loci from these hosts, the point estimate for divergence time produces an average third-site *Wolbachia* substitution rate of 2.2×10^−9^ substitutions/3rd position site/year, identical to the *Nasonia* estimate. Using the 95% credible interval for the host divergence times from Suvorov *et al*. (2022), a 95% credible interval for the substitution rate is (1.9–2.6)×10^−9^ substitutions/third-position site/year. Our analysis of 613,605 bp of *Wolbachia* data from *Nomada* (see Table 2) indicates a lower average substitution rate per third-position site per year of roughly 5.6×10^−10^ (with 95% credible range [0.40–1.16]×10^−9^). We consider four alternative priors for the *Wolbachia* third-position substitution-rate based on these estimates.

Given the approximations involved in these substitution-rate estimates, it is informative to evaluate the robustness of our results to alternative priors. We consider alternative priors (Table S5) that attempt to capture in different ways the variation seen in Table 1. Rather than try to approximate the asymmetrical confidence intervals for *Nomada* and *Drosophila* with gamma distributions, we used priors that explored different approximations of the variation of our substitution-rate estimates. All four priors assume a mean substitution rate of 1.65×10^−9^ per third-site per year for the *Wolbachia* genomes (this approximation gives two-thirds weight to the concordant estimates from *Nasonia* and *Drosophila* and one-third weight to the *Nomada*-based estimate). Two of our priors assume unimodal distributions of the substitution rates, with variance that approximates the differences between the estimates from *Nomada* versus *Nasonia* and *Drosophila*, and two assume bimodal distributions, with one mode (given two-thirds weight) corresponding to the *Nasonia* and *Drosophila* mean and the other mode (with one-third weight) corresponding to the *Nomada* mean. To explore the consequences of different shapes and variances for the prior distributions of substitution rates, we use both normal and uniform distributions.

Our heuristic approach to describing substitution-rate variation is consistent with the many approximations that enter the estimates of host divergence times. Our informal credible intervals are based on the credible intervals for the host divergence times. Our four substitution-rate priors will be denoted N1, N2, U1 and U2. Let N(1, s) denote a normal random variable with mean 1 and standard deviation s, and let U(a, b) denote a uniform distribution over the interval (a, b). N1 samples rates from N(1, 0.34)×1.65×10^−9^, U1 samples from U(0.33,1.67)×1.65×10^−9^, N2 samples from N(1, 0.36)×0.56×10^−9^ with probability 1/3, and from N(1, 0.08)×2.2×10^−9^ with probability 2/3, and U2 samples from U(0.3,1.7) ×0.56×10^−9^ with probability 1/3, and from U(0.84,1.16)×2.2×10^−9^ with probability 2/3. Table S5 shows how node age estimates and support intervals vary with the alternative priors. The point estimates are quite robust, as expected given the common mean rate for all four priors. The chronograms in Fig. 1 use prior N1.

### Detecting recombination and its influence on estimated phylograms and chronograms

Prior work using only a few genes has identified a pattern of recombination between relatively diverged *Wolbachia* variants (Werren and Bartos 2001; Jiggins *et al*. 2001; Baldo *et al*. 2006). To test for recombination across the *Wolbachia* genome, we used a genetic algorithm (GARD) (Kosakovsky Pond *et al*. 2006), plus three other statistical methods implemented in PhiPack (Bruen *et al*. 2006). We focused our analyses on single-copy genes that were at least 300 bp in length, with minimum recombination segments of 100 bp. We set GARD to detect a maximum of two potential breakpoints. We first completed these analyses using our 20 *w*Mel-like variants to determine the extent of recombination between these closely related *Wolbachia*. 411 genes met our criteria for this analysis.

To test for recombination between more distantly related *Wolbachia*, we completed a second analysis using *w*Mel and *w*Ri, plus three other A-group strains (*Wolbachia* associated with *Andrena hattorfiana*, *Anoplius nigerrimus*, and *Apoderus coryli*), and B-group *w*Mau in *Drosophila mauritiana* (Meany *et al*. 2019; Vancaester and Blaxter 2023). We searched for homologs of all 1292 genes in *w*Mel in these five other *Wolbachia*. Homologs for 1124 genes were found in all 5 *Wolbachia*, of which 111 were shorter than 300pb and excluded. The remaining 1013 were tested for recombination using GARD and the three statistical tests implemented in PhiPack. Because recombination could potentially influence our estimation of phylograms and chronograms, we also revisited the phylogram and chronogram analyses presented in Meany *et al*. (2019) that included 9 group-A and 6 group-B strains. We identified genes with no evidence of recombination according to all 4 tests described above and used them to re-estimate a Bayesian phylogram and an absolute chronogram with RevBayes 1.1.1, as in Meany *et al*. (2019). We estimated the phylogram with the GTR + Γ model, partitioning by codon position (*5*). To estimate the absolute chronogram, we first estimated a relative relaxed-clock chronogram with the root age fixed to 1, partitioned by codon position. The relaxed-clock branch-rate prior was Γ(7,7), normalized to a mean of 1 across all branches. We transformed the relative chronogram into an absolute chronogram using both the original and new priors for *Wolbachia* 3rd position site/year substitution rates discussed above.

### Host phylogeny and chronograms

A key conclusion of our analyses is that closely related *Wolbachia* are transferred among distantly related hosts on a time scale many orders of magnitude faster than host divergence times. Our estimates concerning host divergence rely on recent calibrations from the literature. Fig. 1B presents our most diverged hosts. The clades Diptera and Hymenoptera span the Holometabola. According to the fossil-calibrated chronograms in Wang et al. (2016, their Fig. 3), Diptera and Hymenoptera diverged ∼350 million years ago (MYA), with 95% highest posterior density credibility interval (HPD CI) of (378–329 MYA, Devonian–Carboniferous). This is consistent with the point estimates produced by Misof *et al*. (2014) and Johnson *et al*. (2018) (Fig. 1). The placement of the family Diopsidea, the stalk-eyed fly clade that includes *Sphyracephala bevicornis*, within superfamily Diopsoidea in the paraphyletic acalyptrate group of Schizophora remains uncertain (Bayless *et al*. 2021). Hence, the maximum divergence time between *Sphyracephala bevicornis* and any drosophilid is the crown age of the Schizophora, which includes both the Drosophilidae and Diopsoidea. The minimum divergence time between the Drosophilidae and Diopsoidea is the crown age of the Drosophilidae (which certainly excludes the Diopsoidea). Wiegmann *et al*. (2011) (Fig. 3) estimate the crown age of the Schizophora at ∼70 MYA. Suvorov *et al*. (2022) (Fig. 1) estimate the crown age of the Drosophilidae at about 47 MYA (with 95% HPD CI of 43.9–49.9, Devonian–Carboniferous). Our approximate point estimate in Fig. 1B for the divergence of Drosophilidae and Diopsoidea, 59 MY, is the midpoint of these bounds.

For the drosophilids in Fig. 1C, node ages and approximate confidence intervals were estimated from the fossil-calibrated chronogram of Suvorov *et al*. (2022), using supplementary information as needed. We number the 12 nodes in Fig. 1C from left to right, with 1 denoting the divergence between the subgenera *Sophophora* (including *D. tropicalis*) and *Drosophila* (including *D. arawakana*), 2 denoting the divergence of *D. tropicalis* from the *D. melanogaster* subgroup, …, 11 denoting the crown age of the *D. melanogaster* subgroup, and 12 denoting divergence time between *D. borealis* and *D. incompta*. For species in our Fig. 1C that are not included in Fig. 1 of Suvorov *et al*. (2022), we used the NCBI Taxonomy Browser to determine the closest relative(s) included in Suvorov *et al*. (2022). We estimated node ages and approximate CIs from the x-axis of their Fig. 1, using the measurement tool in Adobe Acrobat Pro DC (ver. 2022.001.20169), converting distances to time using the scale bar at the bottom of their figure. Our symmetrical approximate CIs were obtained by measuring the widths of the CI profiles in Fig. 1 of Suvorov *et al*. (2022) (*21*). This method produced the approximate ages and CIs for nodes 1– 11 in our Fig. 1C. As a check, our approximation method produces 47 ± 2.8 MY as the crown age in Fig. 1C; in their text, Suvorov *et al*. (2022) estimate this age as ∼47 MYA with 95% CI 43.9–49.9 MYA.

We used our Suvorov *et al*. (2022) calibrations to set the crown ages of the *melanogaster* and *montium* subgroups (nodes 10 and 11). Within those subgroups and for node 12 (*D. borealis*, *D. incompta*), we estimated relative divergence using relaxed-clock relative chronograms under a GTR + Γ [7,7] model of molecular evolution, following the methods in Conner *et al*. (2021), summarized below. Estimating the divergence time between *D. borealis* and *D. incompta* was the most problematic. *D. incompta* belongs to the *D. flavipilosa* species group (De Ré *et al*. 2017). Both nuclear and mtDNA data indicate that among the host species we analyzed, *D. incompta* is most closely related to *D. borealis* in the *D*. *virilis* species group.

To estimate relative divergence times for host drosophilids not included in Suvorov *et al*. (2022) (*21*), we obtained genomes from NCBI (Table S1). Coding sequences for the 20 nuclear genes used in the analyses of Turelli *et al*. (2018) (*aconitase, aldolase, bicoid, ebony, Enolase, esc, g6pdh, GlyP, GlyS, ninaE, pepck, Pgi, Pgm1, pic, ptc, Tpi, Transaldolase, white, wingless,* and *yellow*) were obtained from FlyBase for *D. melanogaster*. We used tBLASTn to identify orthologs in the other genome assemblies. The sequences were aligned with MAFFT v. 7 (*4*) and trimmed of introns using the *D. melanogaster* sequences as a guide. We estimated a relaxed-clock relative chronogram with RevBayes 1.1.1 (*5*) with the root age fixed to 1 using the GTR + Γ [7,7] model, partitioned by gene and codon position. We used the same birth-death prior as Turelli *et al*. (2018) (*6*). Each partition had an independent rate multiplier with prior Γ(1,1), as well as stationary frequencies and exchangeability rates drawn from flat, symmetrical Dirichlet distributions. The branch-rate prior was Γ(7,7), normalized to a mean of 1 across all branches. Four independent runs were performed, which agreed with each other. Nodes with posterior probability less than 0.95 were collapsed into polytomies.

### CI assays

To test for cytoplasmic incompatibility (CI) in *D. seguyi* and *D. bocqueti*, we estimated the egg hatch frequencies from putatively incompatible crosses between females without *Wolbachia* and males with *Wolbachia* (denoted IC) and the reciprocal compatible cross (CC) between females with and males without *Wolbachia*. With CI, we expect lower egg hatch from IC crosses than from CC crosses. To generate lines of both species without *Wolbachia*, we exposed *Wolbachia*-carrying lines to tetracycline-supplemented (0.03%) cornmeal medium (see Shropshire *et al*. 2021 for details) for three generations. We confirmed the absence of *Wolbachia* in the treated flies using PCR within two generations of tetracycline treatment using primers for the *Wolbachia*-specific *wsp* gene (Braig *et al*. 1998; Baldo *et al*. 2005) and a second reaction for the arthropod-specific 28S rDNA as a host control (Nice *et al*. 2009). Our PCR thermal profile began with 3 min at 94C, followed by 34 rounds of 30 sec at 94C, 30 sec at 55C, and 1 min and 15 sec at 72C. The profile finished with one round of 8 min at 72C. We visualized PCR products using 1% agarose gels that included a molecular weight ladder. The stocks were maintained and experiments were conducted in an incubator at 25°C.

We reciprocally crossed *Wolbachia*-carrying *D. seguyi* and *D. bocqueti* lines to their tetracycline-treated conspecifics. Tetracycline-treated stocks were given at least four generations to recover prior to our experiments. Virgins were collected from each line and placed into holding vials for 48 hr. We set up each IC and CC cross with one female and one male in a vial containing a small spoon with cornmeal medium and yeast paste for 24 hr. Males and females were two days old at the beginning of these experiments. Each pair was transferred to a fresh vial every 24 hr for 5 days. We counted the number of eggs laid at the time that adults were transferred to new vials and the number of eggs that hatched were scored after an additional 24 hrs. The data analyzed were hatch proportions for crosses across the 5-day period. To control for cases where females may not have been inseminated, we excluded crosses that produced fewer than 10 eggs. We used one-sided Wilcoxon tests to determine whether IC crosses produce lower egg hatch proportions than do CC crosses.

### Extracting *cif* sequences

We used BLAST to identify contigs with *cif* sequences, using *cif_w_*_Mel[T1]_, *cif_w_*_Ri[T2]_, *cif_w_*_No[T3]_, *cif_w_*_Pip[T4]_, *cif_w_*_Stri[T5]_, *cif_w_*_Tri[T5]_, and *cif_w_*_Bor[T5]_ as query sequences. We used Genious Prime to extract open-reading frames with blast homology to *cif* sequences for downstream analyses (Kearse *et al*. 2012). Among all *cif* sequences, only *cifB_wMal[T1]_* did not have a clear associated ORF; however, we did observe sequence homology in the region. We assigned the *cif* sequences to Types (T1–T5) based on similarity to reference genes of each Type. Table 3 provides the sources of all *Wolbachia* genomes used in our *cif* sequence analyses.

### Extracting serine recombinase genes

We used the large sr to categorize *Woviruses* as sr1WO, sr2WO, sr3WO, or sr4WO (Bordenstein and Bordenstein 2022). We used WOCauB3, WOVitA1, WOMelB, and WOFol2 sr as queries in BLAST searches. If the *Wolbachia* assembly clearly assigned an sr sequence to a phage, we assigned the phage to sr1WO, sr2WO, sr3WO or sr4WO based on the similarity to the reference sequences.

### Phylogenetic topological identity

We tested for discordance between phylogenetic trees estimated from different data, e.g., *cifA* versus sr sequences within the same *Woviruses* using the SH (Shimodaira and Hasegawa 1999) and AU (Shimodaira 2002) tests, as implemented in IQ-Tree (Minh *et al*. 2020). Unlike the topological-similarity tests described below, the null hypothesis in these tests is topological identity.

### Phylogenetic topological similarity

We used normalized Clustering Information (CID), Jaccard-Robinson-Foulds (JRF), and Robinson-Foulds (RF) distances to test for similarity between pairs of trees, as described by Smith (2020). All metrics were calculated using the TreeDist package in R (Smith *et al*. 2023). To normalize the distance metrics, we divided the observed value by the mean distance obtained from comparing 10,000 random tree pairs with equal numbers of leaves. We generated random trees using the *ape* package in R (Paradis *et al*. 2023). We calculated *P*-values as the proportion of random trees of the same size as our data that produced distances below the observed distance. This tests the hypothesis that two trees are more similar than expected by chance.

### Characterizing Cif protein structures

We ran HHPred on a Linux kernel to identify putative functional domains using the Pfam-A_v35 and SCOPe70_2.07 databases (Zimmermann *et al*. 2018). We considered only annotations with > 80% probability. If alternative annotations produced probability > 0.8, we selected the annotation with the highest probability. We used AlphaFold2 (Jumper *et al*. 2021) to predict Cif structures. Entries in the “reduced database” provided with AlphaFold prior to 5/10/22 were used to generate multiple sequence alignments (MSA) within AlphaFold. We generated five structures for each protein and performed amber relaxation to prevent unrealistic folding patterns. We sorted the five relaxed models by mean pLDDT and used the top result in other analyses. We visualized protein structures using PyMol 2.5.2 (Schrödinger, LLC 2015) and aligned proteins relative to Cif*_w_*_Mel[T1]_ for imaging using Cealign.

We generated TM-scores in PyMol used for pairwise-comparisons of Cif proteins to determine Cif structural similarities using the psico module (Holder and Schmidt 2023). We performed each analysis twice, switching the reference and target trees. This impacts the TM-score because the score is normalized to the length of the target protein. We used a Mann-Whitney U test in R to compare the TM-scores from CifA and CifB—truncated proteins and proteins at the edge of a contig were removed from this analysis. We assessed the relationship between TM-score and percent identity using a Spearman correlation.

### Characterizing Cif selective pressures

To identify evidence of selection along the Cif proteins, we calculated the ratio of the number of non-synonymous substitutions per non-synonymous site (*d_N_*) to the number of synonymous substitutions per synonymous site (*d_S_*) using the Sliding Window Analysis of Ka and Ks (SWAKK) webserver (Liang *et al*. 2006). SWAKK calculates *d_N_*/*d_S_* by generating an alignment of two nucleotide sequences, mapping the alignment onto the tertiary structure, and calculating *d_N_*/*d_S_* with a 10 Å spherical sliding window across the reference structure. We used *cif_wMel_*_[T1]_, *cif_wRi_*_[T2]_, *cif_wApo_*_[T3]_, *cif_wTei_*_[T4]_, and *cifA_wTri_*_[T5]_ as references for each *cif* Type. We used AlphaFold structures for tertiary mapping. All statistical analyses were performed using the median *d_N_*/*d_S_* for each site across pairwise comparisons. We calculated BCa 95% confidence intervals for *d_N_*/*d_S_* values using the boot package in R (Canty and Ripley 2024).

### Figure generation

We produced and/or edited figures in R, Figtree, Inkscape 1.1, Adobe Illustrator, and Keynote.

## Data availability

Source data for the main and Supplementary Data figures are provided in the online version of this paper or in Dryad. Newly sequenced *Wolbachia* genomes are available under BioProject PRJNA1021588.

## Author Contributions

Conceptualization: JDS, MT, BSC; Data curation: WRC, DV; Formal analysis: JDS, WRC, DV, MT, BSC; Funding acquisition: JDS, BSC; Project administration: JDS, MT, BSC; Resources: BSC; Supervision: MT, BSC; Validation: JDS, WRC, DV; Visualization: JDS, DV, BSC; Writing – Original draft: JDS, MT, BSC; Writing – Review & editing: JDS, WRC, DV, AAH, MT, BSC.

## Acknowledgments

We thank Daniel Matute for directing us to data for *Zaprionus* species. Nitin Ravikanthachari and John Statz for comments that improved the manuscript and Tim Wheeler for support in the laboratory.

