## Supporting Information for "Rapid host switching of *Wolbachia* and even more rapid turnover of their phages and incompatibility-causing loci"

#### Supplementary Discussion

##### *Recombination*

We detected little evidence for recombination among our 20 *w*Mel-like *Wolbachia* presented in Fig. 1A (GARD: 0/411 genes, PHI test: 0/411 = 0%, Max $\chi^2$ test: 3/411 = 0.7%, Neighbour Similarity Score: 0/411 = 0%). To test for recombination among more diverged variants, we completed a separate analysis that included *w*Mel and *w*Ri, three other group-A variants (*Wolbachia* associated with *Andrena hattorfiana*, *Anoplius nigerrimus*, and *Apoderus coryli*), and B-group *w*Mau (Meany *et al.* 2019; Vancaester and Blaxter 2023). Consistent with previous analyses of recombination among distantly related *Wolbachia* variants (Werren and Bartos 2001; Jiggins *et al.* 2001; Baldo *et al.* 2006), GARD found evidence of recombination in 728 genes, with recombination in more than half of these genes also supported by the tests implemented in PhiPack (PHI: 418/728, Max $\chi^2$: 485/728, NSS: 520/728). Recombination was detected by all three tests in 353 genes. These 353 genes include 245 with a gene region where *w*Mel and *w*Ri are sister. Of these, 7 have a region where *w*Mel and *w*Mau are sister, 25 have a region where *w*Ri and *w*Mau are sister, 37 have a region where *w*Mel and *A. nigerrimus Wolbachia* are sister, and 38 have a region where *w*Ri and *A. nigerrimus Wolbachia* are sister. Our new analyses generalize prior results that used only a few genes and found support for pervasive recombination between relatively diverged *Wolbachia* (Werren and Bartos 2001; Jiggins *et al.* 2001; Baldo *et al.* 2006), while confirming that our criteria for selecting genes to produce phylograms indirectly selects for genes with little evidence for recombination.

Recombination between group-A and group-B *Wolbachia* could potentially influence estimated phylograms and chronograms. To test this hypothesis, we searched for genes with no evidence of recombination using the 15 *Wolbachia* we previously analyzed (Meany *et al.* 2019). Of 168 single-copy and equal-length genes, we discarded 14 that were shorter than 300 bp. All three statistical tests found no evidence for recombination in 51 of the remaining 154 genes. We used these 51 genes to re-estimate a phylogram and absolute chronogram with RevBayes 1.1.1. Four independent runs of each were performed and all agreed. With the exception of the placement of *w*AlbB (Fig. S1A), our phylogram estimated using only the 51 genes with no evidence for recombination recapitulates the topology estimated by Meany *et al.* (2019) (Fig. S1B). Our estimate of A-B group divergence time using only these 51 genes (~7-38 MYA) also overlaps with our previous relaxed-clock analyses with branch-rate priors $\Gamma(7,7)$ (~8-46 MYA) and $\Gamma(2,2)$ (~6-36 MYA), as well as with our strict-clock analysis (~12-64 MYA) (Meany *et al.* 2019). As expected, using our new calibration the estimated divergence times are longer for both the 51 gene set (~32-165 MYA) and the original Meany *et al.* (2019) gene set (~42-196 MYA); but again divergence estimates are overlapping. This confirms that our previously estimated phylograms and chronograms are insensitive to the inclusion of genes with evidence of recombination.

##### *Inferences from chronograms*

Inferences from these chronograms are limited with only one available *Wolbachia* sequence from most of these hosts. For instance, we cannot determine which *Wolbachia* infection was acquired first, whether two sister tips on our *Wolbachia* chronograms correspond to one of those hosts acquiring its *Wolbachia* (by either introgression or non-sexual horizontal transmission) from the host of the sister *Wolbachia* variant – or whether one or both *Wolbachia* were recently acquired from an unsampled intermediate host. Turelli *et al.* (2018) previously proposed that *w*Ri in *D. simulans* was acquired horizontally from *D. ananassae*, because multiple *Wolbachia* sequences were available from each species, and the *w*Ri sequences formed a clade nested within a paraphyletic cluster containing the *Wolbachia* from *D. ananassae*. As noted by Richardson *et al.* (2012), divergence times for extant *Wolbachia* variants in a host species may be much shorter than the duration of its current *Wolbachia* variant, because selective sweeps and genetic drift will eliminate sequence variation that indicates the duration of associations.

Fig. S2. illustrates the limited inferences about *Wolbachia* divergence and acquisition possible from individual *Wolbachia* sequences from various hosts. We consider four hypothetical host species with horizontally acquired *Wolbachia*. Host 1 acquires its *Wolbachia*, denoted $w_1$, at time $T_1$, and we denote by $T_i(j » i)$ the time at which host i acquires its *Wolbachia*, denoted $w_i$, from host j. If we postulate $T_1 > T_2(1 » 2) > T_3(1 » 3) > T_4(2 » 4)$, the resulting chronogram is Fig. S2. The chronogram provides no clue that variant $w_1$ was acquired first or that $w_2$ was acquired from host 1. Thus, we can infer only that all but one of the variants shown in Fig. 1A were acquired in the last 263–1.2 MY and that a transfer between a dipteran and a hymenopteran host occurred within the last 184–817 KY, possibly mediated by intermediate hosts. Moreover, the chronogram Fig. 1A is consistent with many of these hosts, for which we do not have multiple *Wolbachia* sequences, acquiring their current *Wolbachia* very recently, followed by spatial spread as observed for *Wolbachia* in several hosts (Turelli *et al.* 2018).

##### *Correction of Turelli et al. (2018) concerning the acquisition of* w*Spc*

The analyses presented in our text deal with only a single *Wolbachia* sequence from each host species. As noted in Fig. S2, single sequences do not allow us to determine the direction or temporal order of *Wolbachia* acquisitions. In contrast, Turelli *et al.* (2018) had multiple *Wolbachia* sequences from several species. Specifically, the *w*Ri sequences from *D. simulans* formed a clade nested within a paraphyletic cluster of *w*Ana sequences from *D. ananassae*. Given that *D. ananassae* and *D. simulans* span the *D. melanogaster* species group, which diverged on the order of 47 MYA (see Fig. 1C), introgression is impossible. Hence we conjectured that horizontal non-sexual transmission of *Wolbachia* occurred between *D. ananassae* and *D. simulans*, with *D. ananassae* as a plausible source. Turelli *et al.* (2018, Fig. 1) also analyzed eight *Wolbachia* sequences from *D. suzukii* (*w*Suz) and one *D. subpulchrella* (*w*Spc). Two copies of *w*Suz from Asia (the other *w*Suz samples came from North and South America) formed a clade with the single *w*Spc, sister to the other six *w*Suz sequences. We used these data to conjecture that *D. suzukii* was a plausible source of the *Wolbachia* in *D. subpulchrella*. However, the two putative Asian *w*Suz sequences, obtained from GenBank, were in fact derived from *D. subpulchrella* specimens (China-AA27 and Korea-AA7). This misidentification was first kindly pointed out to us by Mathieu Gaultier, then confirmed by Joanna Chiu. With this correction, the data in Fig. 1 of Turelli *et al.* (2018) produce sister clades of six *w*Suz and three *w*Spc, providing no information on the direction of transfer, as illustrated in Fig. S2.

##### cif *sequence and Cif structure*

We used up to three pieces of information—strain, type designation, and copy number—in a subscript behind the gene name to describe specific *cifs*. For example, *w*Zta *Wolbachia* encode two T1 *cifA* variants that we denote $cifA_{wZta[T1\text{-}1]}$ and $cifA_{wZta[T1\text{-}2]}$. Cif proteins are diverse (Fig. S3) and subject to putative pseudogenization with early in-frame stop codons (discussed below). Putatively intact CifA proteins range from 344 to 493 aa ($CifA_{wZta[T4]}$–$CifA_{wDal[T1\text{-}2]}$) and CifB proteins are between 546 to 4433 aa ($CifB_{wSan[T4]}$–$CifB_{wSbr[T5]}$) (Fig. S4A). The shortest $CifB_{[T5]}$ ($CifB_{wTri[T5]}$ = 2833 aa) is 1304 aa larger than the largest CifB from other Types ($CifB_{wZta[T1\text{-}2]}$ = 1529 aa). CifA homologs lack confident structural-homology-based domain annotations (Probability > 0.8, see Söding *et al.* 2005) for a description of HHpred homology detection and structure prediction) (Fig. S4A) (Lindsey *et al.* 2018; Martinez *et al.* 2021). Conversely, all CifB homologs encode a PD-(D/E)XK nuclease (Nuc hereafter). $CifB_{[T1]}$ encode a Ulp1 deubiquitilase (Dub hereafter), and $CifB_{[T5]}$ uniquely encode OTUB2 deubiquitilase, R12E2.13-like mammalian-wide interspersed repeat, BurrH DNA-binding, Latrotoxin_C toxin, and tetratricopeptide repeat domains (Fig. S4A).

To investigate Cif protein sequence and structural similarity, we calculated pairwise amino-acid identity (ID) and template-modeling scores (TM) (Zhang and Skolnick 2004). TM-scores indicate the degree of similarity between two superimposed structures, with TM = 0 indicating no similarity, TM = 1 corresponding to complete congruence, and TM > 0.5 indicating highly similar (Xu and Zhang 2010). We found that putatively intact $CifA_{[T1]}$ proteins are significantly more similar than $CifB_{[T1]}$ from the same pairs ($N$ = 18) in terms of both sequence (95% BCa confidence intervals: $ID_{CifA[T1]}$ = 0.78 – 0.86, $ID_{CifB}$ = 0.69–0.79; $P$ = 0.0002; Fig S3A) and AlphaFold structure ($TM_{CifA}$ = 0.89 – 0.92, $TM_{CifB}$ = 0.56–0.62, $P < 10^{-10}$; Fig. 3B). We observed similar trends across Cif Types ($N$ = 28 pairs; Fig. S3A) and among 12 $Cif_{[T1]}$ from *w*Mel variants and six from *w*Ri variants (Fig. S3B and C).

##### **Wovirus** *rapidly turnover among* **Wolbachia** *genomes*

We characterized the diversity and distribution of sr alleles in closely and more distantly related *Wolbachia* to understand *Wovirus* association with these *Wolbachia*. sr1WO occurs in all *w*Ri-like *Wolbachia* except for *w*Aur and *w*Tri, while sr2WO occurs in all *w*Ri-like variants (except *w*Tri), *w*Mel-like *w*Tris, and the four members of the *w*SYTZ clade. sr3WO is the most commonly observed *Wovirus*, present in one-to-three copies in all *w*Mel-like and *w*Ri-like *Wolbachia* genomes (Table S3). In some cases we observe congruent *Wolbachia*-*Wovirus* topologies. For example, sr1WO exhibits perfect phylogenetic congruence with *w*Ri-like *Wolbachia* (Fig 1C), while sr2WO (*CID* = 0.49, *P* = 0.0004; *JRF* = 0.50, *P* = 0.04; *RF* = 0.67, *P* = 0.0003) and sr3WO exhibit imperfect (but non-random) congruence with their respective hosts (*CID* = 0.73, *P* = 0.0; *JRF* = 0.72, *P* = 0.12; *RF* = 0.94, *P* = 0.0) (Fig S5A and B). We describe the distribution below. Multiple observations indicate *Wovirus* turnover among *Wolbachia* genomes, which we highlight for sr2WO and sr3WO *Wovirus* below.

First, sr2WO *Wovirus* form two clades: a clade observed in all *w*Ri-like *Wolbachia*, and a clade observed in the *w*Mel-like *w*SYTZ *Wolbachia* clade (MRCA 54–353 KYA) and in distantly related *w*Mel-like *w*Tris. While the history of sr2WO cannot be fully resolved from these data, we hypothesize that sr2WO in the *w*Ri-like clade, the *w*SYTZ clade, and *w*Tris represent independent acquisition events that occurred after *w*Mel-like and *w*Ri-like *Wolbachia* divergence

(MRCA 263–1.2MY). The absence of sr2WO in *w*Rec—sister to *w*Tris—implies gain or loss of srWO2 may have occurred as recently as 101–762 KYA. Figure S5A presents these patterns.

Second, each *w*Ri-like strain also contains closely related sr3WO *Wovirus*, with duplications observed in in *w*Ri, *w*Ana, and *w*Suz *Wolbachia* genomes. These sr3WO *Wovirus* observed in *w*Ri-like *Wolbachia* form a clade that is sister to an sr3WO clade observed in several *w*Mel-like variants: the four *w*SYTZ strains; sister (*w*Seg,*w*Mal); a clade of three strains (*w*Sbr, (*w*Ara, wSpa)); as well as *w*Bocq, *w*Bor, and *w*Dal. The absence of this haplotype in *w*Ach indicates gain or loss since its divergence from *w*Bocq in the last 14–106KY. Similarly, absence from *w*Au, and *w*Tro indicates gain and/or loss since their divergence from *w*Bor in the last 179–790KY. We observe an additional sr3WO haplotype that is absent from *w*Bor but present in its 7 closest relatives, implying loss of this haplotype in the last 179–790KY. Fig. S5B presents these patterns, including our observation of another closely related copy of sr3WO *Wovirus* present in all *w*Mel-like *Wolbachia* (MRCA 263–1.2 MYA).

##### cif *turnover among phages*

Cooper *et al.* (2019) found that a single sr3WO *Wovirus* in *w*STY contain both Type 1 and Type IV *cif* operons, whereas closely related *w*Mel does not contain a Type IV *cif*. This discovery has been confirmed with more complete *w*STY assemblies (Baião *et al.* 2021). Thus *cifs* clearly move in and out of *Woviruses,* precluding concordant *cif* and *Wovirus* phylogenies. We used our estimated phylogenies for sr3WO haplotypes (sr3 alleles) and *cifA*$_{[T1]}$ alleles from Fig. 4 to assess how commonly *cifs* move among *Woviruses*. Horizontal *cif* movement among phages will produce discordance between the phylogenies estimated from sr3 alleles versus *cifs*. The SH and AU tests assess discordance by determining whether data from one set of markers is compatible with the phylogeny estimated from the other. As shown in Fig. 3, there is clear discordance in the topologies of *cifA*$_{[T1]}$ and sr3WO among the *w*Mel-like variants. The SH and AU tests assign $P < 10^{-6}$ to the fit of the more resolved sr3WO data to the less resolved *cifA*$_{[T1]}$ tree in Fig. 3.

##### *Origin and movement of* cif *operons in* w*SYT*

The primary mode of *cif* movement is difficult to ascertain without a comprehensive comparative analysis of fully contiguous genomes and associated plasmids. However due to *cifs'* imperfect association with phages, and their proximity to IS elements, *cif* horizontal transfer may plausibly occur via phage-mediated and phage-independent mechanisms. Cooper *et al.* (2019) hypothesized that the Type IV *cifs* in *w*Yak (and more broadly, *w*SYT) could have entered a phage that already carried Type I *cifs* via homologous recombination with extrachromosomal DNA excised by IS elements. This proposal was disputed by Baião *et al.* (2021). Using better assemblies, they proposed that phage probably delivered Type IV *cifs* into the ancestor of the *w*SYT clade and *w*Mel that were then subsequently deleted in *w*Mel. Their proposal is based on the phylogenetic relationship of *octomom* WD0513 homologs. Their hypothesis is implausible for several reasons that become obvious when assessing their assemblies.

First, the Type IV *cifs* in *w*SYT and divergent B-group *w*Pip are < 3% diverged, while the surrounding phage genes in *w*Yak and *w*Pip are ~15% diverged (see Cooper et al. 2019, Figure 5). This points to a common donor for the Type IV *cifs* in *w*Yak and *w*Pip that is distinct from the phages that currently contain them. The highly diverged (~15%) phage sequences between *w*Yak and *w*Mel following the *cifAB* genes Cooper *et al.* (2019, Fig. 5) clearly indicate that this transfer did not occur in the ancestor of *w*SYTM followed by deletion from *w*Mel, as proposed by Baião *et al.* (2021). Baião *et al.* (2021) note that the divergence of the WD0513 gene between *w*Mel and *w*SYT is "much higher than most other parts of the genomes” (~15%) but still conclude that this Octomom associated gene is orthologous to the copy in *w*SYT. This proposal is unreasonable, as most of the *w*Mel and *w*SYT genomes are <1% diverged. Given that the 15% divergence for WD0513 between *w*Mel and *w*SYT is comparable to the divergence for the phage-associated genes flanking the Type IV *cifs* in *w*SYT, it is most plausible that WD0513 was transferred with these phage genes and the Type IV *cifs* from an unknown donor. Incidentally, WD0513 is immediately flanked by a transposon Baião *et al.* (2021, Fig. 2A) which, once again, makes it difficult to ascertain whether the mechanism providing extrachromosomal DNA for homologous recombination is phage-, transposon-, or plasmid-mediated. Our finding of discordance between sr3 and associated *cifA*$_{[T1]}$ phylograms (Fig. 3) generalizes our initial discovery of phage-independent *cif* transfer (Cooper *et al.* 2019).

### Supplementary Figures

#### Figure S1

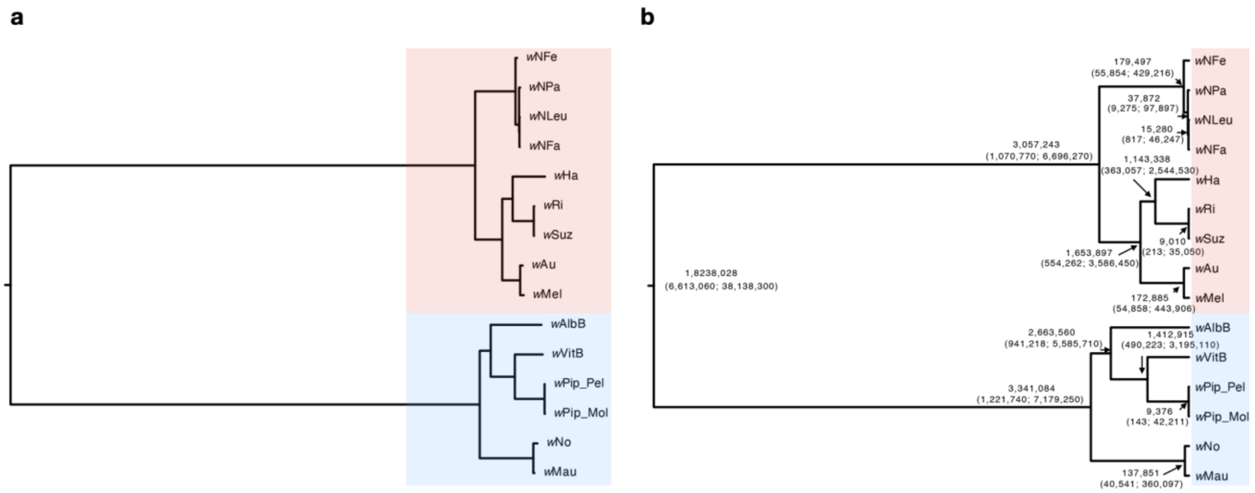


**Fig. S1. Recombination does not influence estimated relationships or divergence times**. **(A)** An estimated phylogram for group-A (red) and group-B (blue) *Wolbachia* strains using only the 51 genes with no evidence of recombination. For this analysis, we follow the approach of Meany *et al.* (2019) as described in our Supplementary Methods. In the main text we also report the divergence of group-A and B *Wolbachia* estimated using our new calibration. Long branches separate the group-A and B clades, with significant variation in the substitution rates across branches. The topology here, with the exception of *w*AlbB placement, fully overlaps with the plylogram presented in Meany *et al.* (2019) that did not consider recombination. Meany *et al.* (2019) placed *w*AlbB outgroup to all other group-B *Wolbachia*. All nodes have Bayesian posterior of one, with the exception of the node distinguishing *w*Nfa and *w*NLeu (0.99) and the node distinguishing *w*AlbB from (*w*VitB,(*w*Pip_Pel, *w*Pip_Mol)) (0.98). Bayesian support less than 1 likely reflects real uncertainty in the placement of these strains. **(B)** An estimated chronogram that includes the same strains. Again, all nodes have Bayesian posterior of 1, with the exception of the node distinguishing *w*AlbB from (*w*VitB,(*w*Pip_Pel, *w*Pip_Mol)) (0.998) and the node distinguishing *w*Nfa and *w*NLeu (0.999). Point estimates for divergence times and their confidence intervals are presented for each node. Estimated divergence times broadly overlap with those estimated by Meany *et al.* (2019) without excluding genes with evidence for recombination.

**Figure S2**

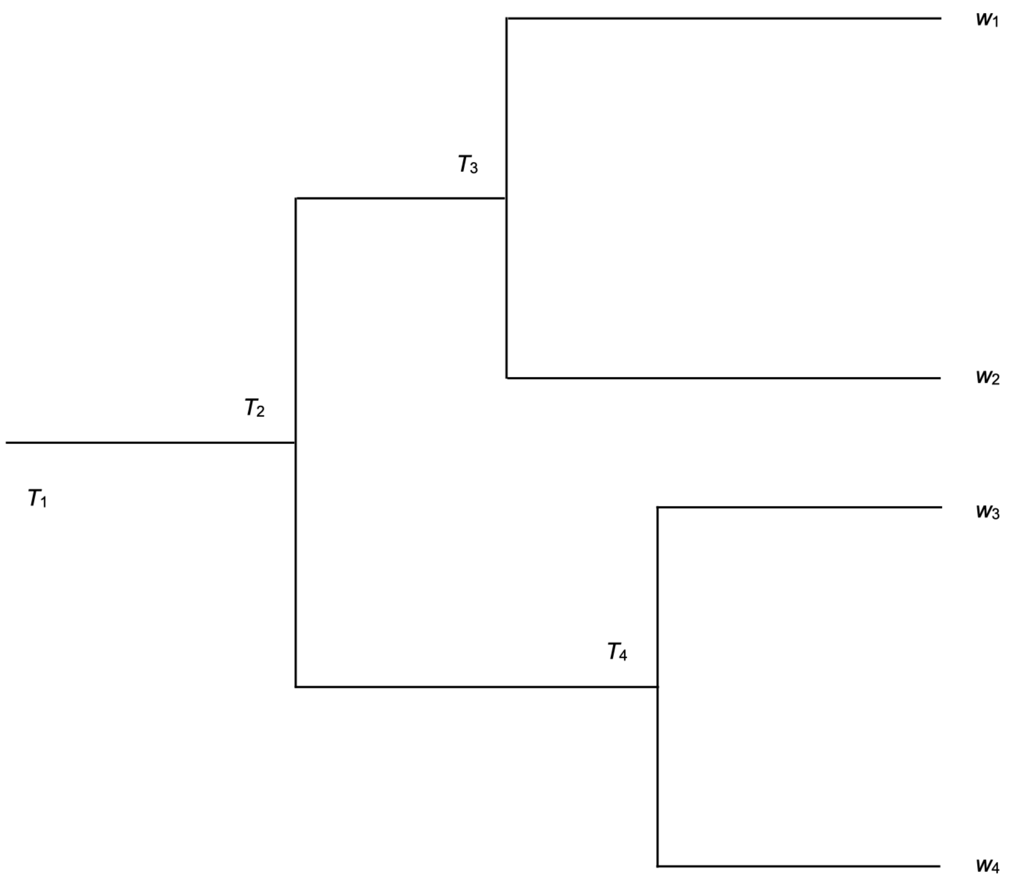


**Fig. S2.** An illustration of the limitations of inferences from the *Wolbachia* chronogram in Fig. 1a (See Supplementary Discussion).

**Figure S3**

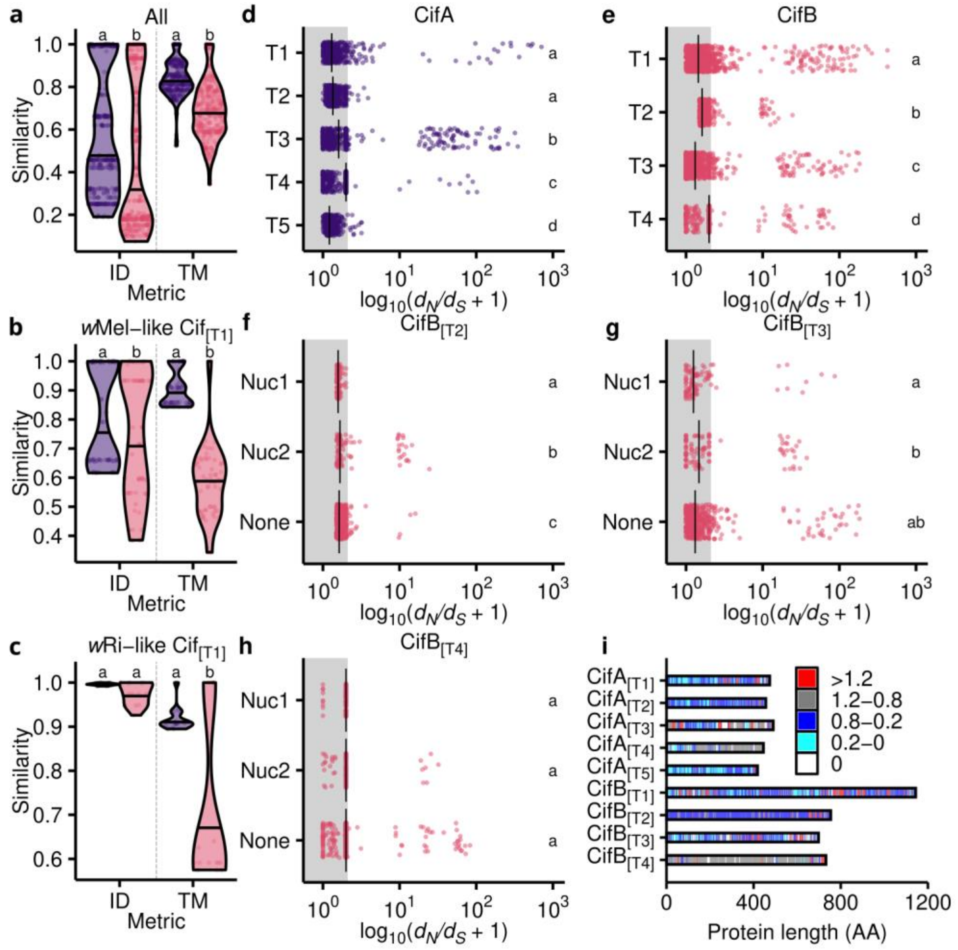


**Fig. S3. Cif protein diversity and selective pressures.** (**A-C**) The relationship between CifA (purple, left) and CifB (pink, right) pairwise sequence identity (ID) and structure similarity (TM). ID and TM are compared across (**a**) intact pairs excluding Cif$_{[T5]}$ ($N = 28$), (**B**) intact Cif$_{[T1]}$ pairs in *w*Mel-like *Wolbachia* ($N = 12$), and (**c**) intact Cif$_{[T1]}$ pairs in *w*Ri-like *Wolbachia* ($N = 6$). (**E-H**) Cif $d_N/d_S$ using a 10 Å spherical sliding window in pairwise comparisons of Cif$_{w\mathrm{Mel}[T1]}$, Cif$_{w\mathrm{Ri}[T2]}$, Cif$_{w\mathrm{Apo}[T3]}$, Cif$_{w\mathrm{Teis}[T4]}$, and CifA$_{w\mathrm{Mel}[T5]}$ to other T1 (CifA: $N = 9$; CifB: $N = 4$), T2 (CifA: $N = 5$; CifB: $N = 2$), T3 (CifA: $N = 5$; CifB: $N = 4$), T4 (CifA: $N = 5$; CifB: $N = 5$), and T5 (CifA: $N = 5$). CifB$_{[T5]}$ was excluded because full-length AlphaFold structures could not be constructed due to computational limitations. (**I**) Median $d_N/d_S$ with a 10 Å window is displayed on linear protein schematics. Shared letters represent statistically similar groups determined by a Mann-Whitney U test (2 groups) or a Kruskal-Wallis and Dunn's multiple comparison test (>2 groups). *P*-values are in Table S7.

**Figure S4**

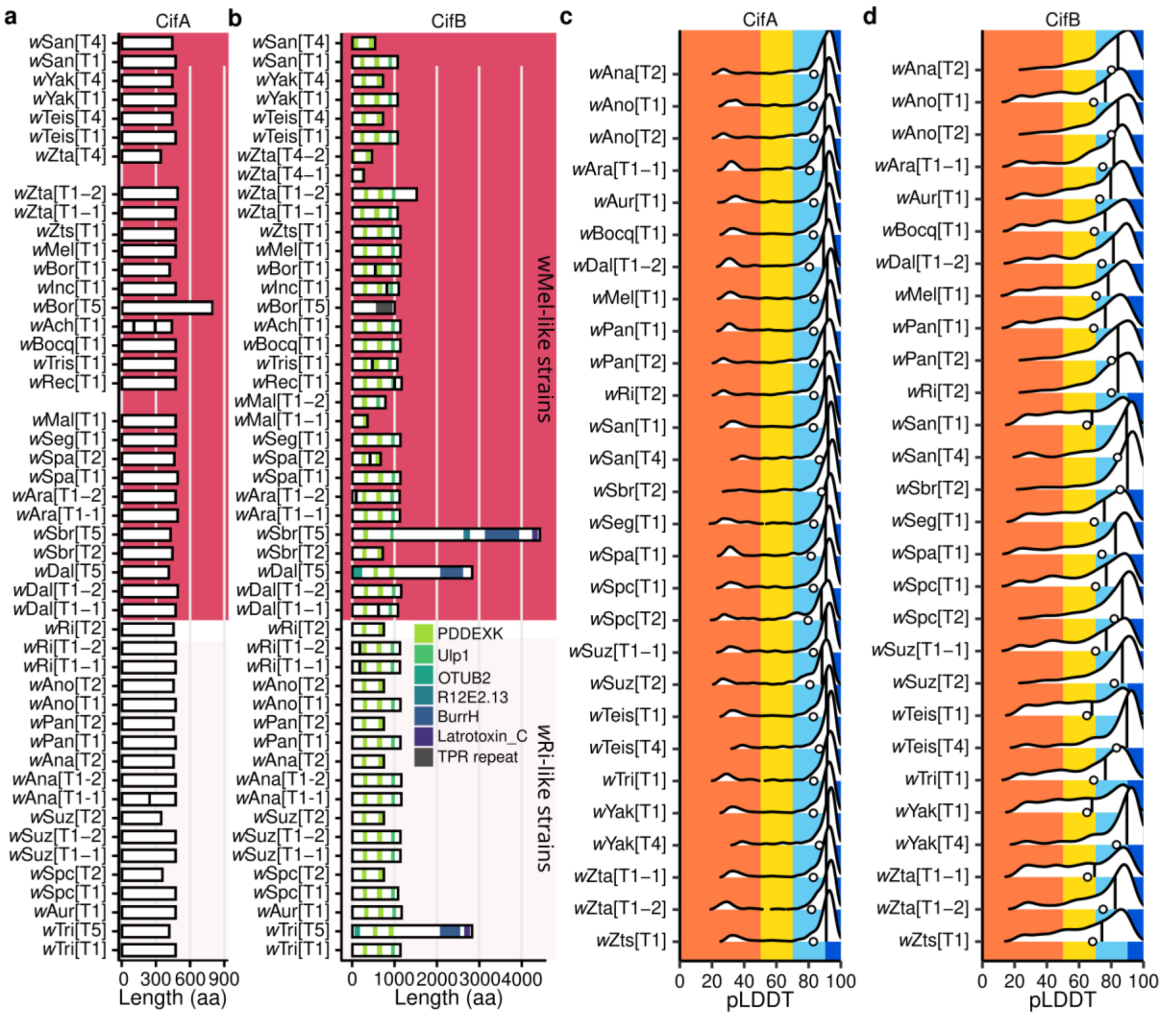


**Fig. S4. Cif domain annotations and sequence/structure diversity.** (**A**) CifA protein schematics. HHPred revealed no annotations with > 80% probability. (**B**) CifB protein schematics showing domains with > 80% probability. Relationship between Cif sequence and structure for Cif$_{[T1]}$ proteins of (**C**) *w*Mel-like and (**D**) *w*Ri-like *Wolbachia*. Structural confidence for (**C**) CifA and (**D**) CifB Alphafold structures. pLDDT is calculated per residue. pLDDT of 100 to 90 (dark blue), 90 to 70 (light blue), 70 to 50 (yellow), and 50 to 0 (orange) indicate very high, high, low, and very low confidence, respectively. Ridgeline plots illustrate the range and density of pLDDT values per protein. Vertical lines and dots represent median and mean pLDDT, respectively.

**Figure S5**

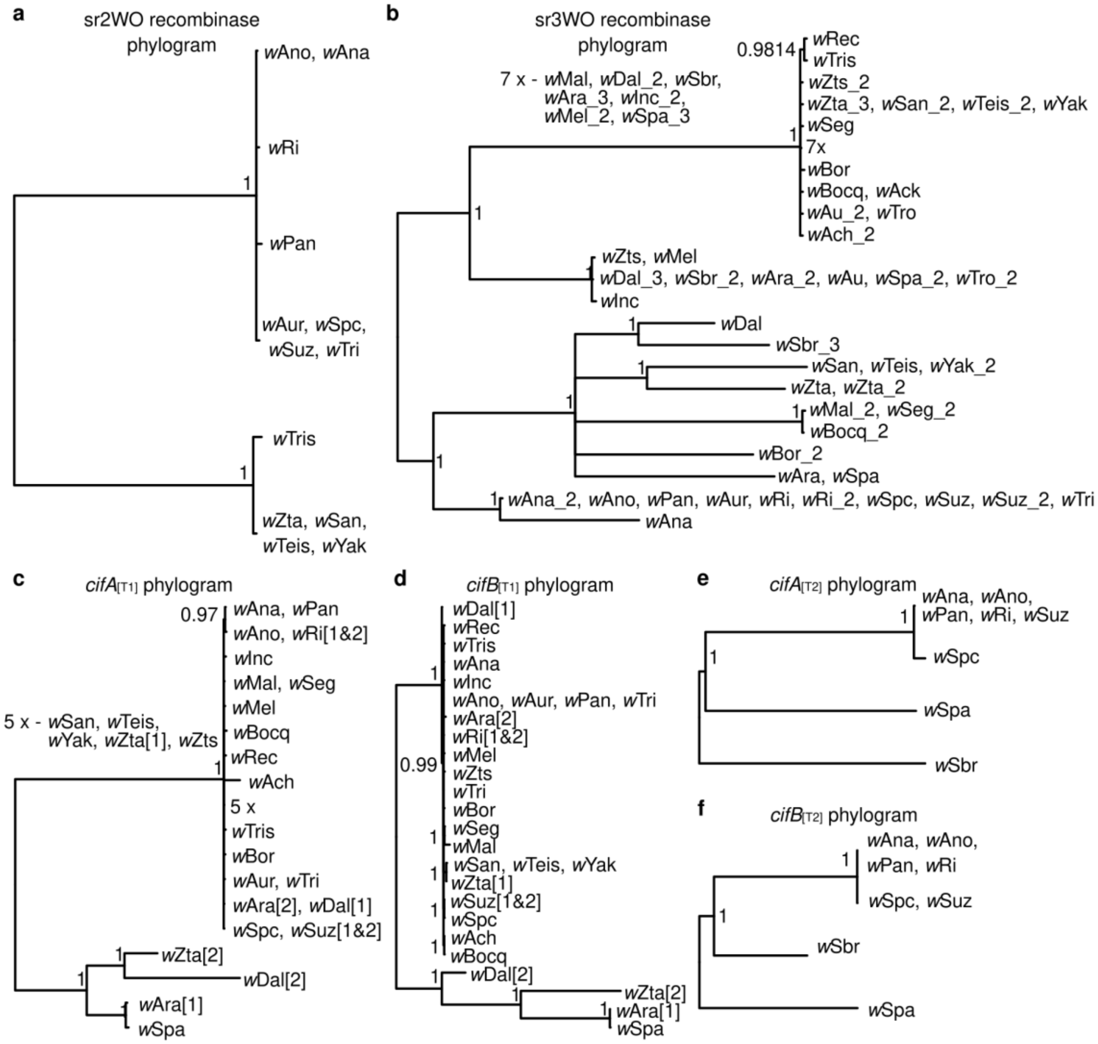


**Fig. S5. Phage and *cif* phylograms.** Serine recombinase phylogenies for homologs of the (**A**) sr2WO phage WOVitA and (**B**) sr3WO phage WOMelB. (**C**) *cifA*[T1], (**D**) *cifB*[T1], (**E**) *cifA*[T2], and (**F**) *cifB*[T2] phylograms with the relaxed clock GTR+G model. All phylograms are midpoint rooted. Node labels represent posterior probability. Identical sequences were collapsed into single tips and nodes with posterior probability < 0.95 were collapsed into polytomies.

#### Supplementary Table Legends

**Table S1.** *Wolbachia* strains, genomes, reproductive phenotypes, and *cifs*.

**Table S2.** Cif sequence and TM score similarity matrices. Pairwise percent identity of all (**A**) CifA and (**B**) CifB proteins reported in this study. Pairwise identity of (**C**) CifA and (**D**) CifB proteins included in the analysis of structure and sequence similarity. (**A-D**) Muscle5 was used to generate 100 multiple sequence alignments (MSA) and the alignment with the highest column confidence was used for downstream analyses. Pairwise identity was calculated as the percent of sites that are shared between pairs in the MSA relative to the length of the pair's alignment, including gaps. TM-scores from pairwise comparisons of (**E**) CifA and (**F**) CifB Alphafold structures.

**Table S3.** Complete information on phage and *cif* typing, and their association with each other and IS elements. **(A)** Summary of the CI phenotypes, *cif* composition, and phage composition for each *Wolbachia* strain in our study. **(B)** Summary of the serine recombinases, their closest *cif*s, and the genes adjacent to the *cifs* in either direction. **(C)** Summary of the *cifs* that we were unable to directly pair with a serine recombinase. We include the associated contig length, and the number of genes on that contig in either direction. **(D)** Summary of serine recombinases without a *cif* gene on the same contig, or where the serine recombinase is further from a *cif* gene than another serine recombinase in the genome. **(E)** Summary of *cifs* with nearby transposases and their identities. **(F)** Summary of the types of transposases found near *cifs* across our genomes, and their distances from focal *cifs*.

**Table S4.** Statistical results associated with main and supporting figures.

**Table S5.** Focal node age estimates and support intervals for the alternative priors described in the Materials and Methods: N1 (normal, unimodal), N2 (normal, bimodal), U1 (uniform, unimodal), and U2 (uniform, bimodal). The point estimates are generally robust, as expected given the common mean rate for all four priors.

**Movie S1**. $d_N/d_S$ mapped on *w*Mel Cif AlphaFold structures. Red, grey, blue, cyan, white, and black represent $d_N/d_S$ values above 1.2, between 1.2 and 0.8, between 0.8 and 0.2, zero, and NA, respectively. The N- and C- termini are displayed with sticks instead of spheres.
